# The Chlamydomonas mitochondrial ribosome: how to build a ribosome from RNA fragments

**DOI:** 10.1101/2021.05.21.445086

**Authors:** Florent Waltz, Thalia Salinas-Giegé, Robert Englmeier, Herrade Meichel, Heddy Soufari, Lauriane Kuhn, Stefan Pfeffer, Friedrich Förster, Benjamin D. Engel, Philippe Giegé, Laurence Drouard, Yaser Hashem

## Abstract

Mitochondria are the powerhouse of eukaryotic cells. They possess their own gene expression machineries where highly divergent and specialized ribosomes, named hereafter mitoribosomes, translate the few essential messenger RNAs still encoded by mitochondrial genomes. Here, we present a biochemical and structural characterization of the mitoribosome in the model green alga *Chlamydomonas reinhardtii*, as well as a functional study of some of its specific components. Single particle cryo-electron microscopy resolves how the Chlamydomonas mitoribosome is assembled from 13 rRNA fragments encoded by separate non-contiguous gene pieces. Novel proteins, mainly helical repeat proteins, including OPR, PPR and mTERF proteins are found in Chlamydomonas mitoribosome, revealing the first structure of an OPR protein in complex with its RNA target. Targeted amiRNA silencing indicated that the novel ribosomal proteins are required for mitoribosome integrity. Finally, we use cryo-electron tomography to show that Chlamydomonas mitoribosomes are attached to the mitochondrial inner membrane via two contact points mediated by Chlamydomonas-specific proteins. Our study expands our understanding of the mitoribosome diversity and the various strategies they adopt for membrane tethering.

**Highlights:**

* Structure of the *Chlamydomonas reinhardtii* mitoribosome
* Fragmented ribosomal RNAs are stabilized by highly intertwined interactions with Chlamydomonas-specific proteins
* Specific r-proteins are essential for rRNA homeostasis and respiratory fitness
* Cryo-ET reveals the mitoribosome association to the inner mitochondrial membrane

## Introduction

Mitochondria are essential organelles of eukaryotic cells that act as metabolic hubs and powerhouses, producing energy through aerobic respiration. They still possess their own genome and gene expression machineries, vestige of their once free-living bacterium ancestor (Eme et al., 2017; Gray, 2012). Due to the evolutionary drift of eukaryotes, mitochondrial complexes involved in metabolism and gene expression combine features from their bacterial ancestor with traits that evolved in eukaryotes (Kummer and Ban, 2021; Waltz and Giegé, 2020). The final step of gene expression, translation, is carried out by specialized mitochondrial ribosomes (mitoribosomes). They synthesize the few proteins still encoded by the mitochondrial genome, most of which are hydrophobic components of the respiratory chain. Despite their shared prokaryotic origin (Martijn et al., 2018), mitoribosome structure and composition were shown to be highly divergent across eukaryotes. They systematically acquired numerous additional ribosomal proteins (r-proteins), and their ribosomal RNAs (rRNAs) were either greatly reduced like in animals and kinetoplastids (Amunts et al., 2015; Greber et al., 2015; Ramrath et al., 2018; Soufari et al., 2020) or expanded like in plants and fungi (Desai et al., 2017; Itoh et al., 2020; Waltz et al., 2019, 2020a).

Among the most prominent unresolved questions in the mitochondria biology field, is the long-standing debate over the peculiar organization of the mitochondrial genome in the unicellular green alga *Chlamydomonas reinhardtii* and the biogenesis of its mitoribosome. This organism is widely used to study photosynthesis and cilia/flagellar motility and function (Harris et al., 2009), but is also an excellent model to investigate mitochondrial biology. It is one of the few organisms where mitochondrial transformation is possible (Remacle et al., 2006), and mitochondrial mutants are viable in photoautotrophic conditions (Salinas et al., 2014). In contrast to vascular plants, or Viridiplantae in general, which are characterized by gene-rich and largely expanded mitochondrial (mt)-genomes, *C. reinhardtii* possess a small linear mt-genome of 16kb. It only encodes eight proteins (all membrane-embedded components of the respiratory chain), three transfer RNAs (tRNAs) and, most intriguingly, non-contiguous pieces of the large subunit (LSU) and small subunit (SSU) ribosomal RNAs (rRNAs), scrambled across the genome (Boer and Gray, 1988; Denovan-Wright and Lee, 1995; Salinas-Giegé et al., 2017). The mt-genome is transcribed as two polycistronic primary transcripts synthesized from opposite strands (Boer and Gray, 1988; Salinas-Giegé et al., 2017). Individual transcripts are then generated from the primary transcripts to produce mature, functional RNAs. Although it was predicted that the rRNA fragments would somehow be integrated into a functional ribosome (Denovan-Wright and Lee, 1995), it is enigmatic how these fragments are recruited, interact with each other, and are stabilized to form the 3D mitoribosome structure.

Here, we combine cryo-electron microscopy (cryo-EM) with *in situ* cryo-electron tomography (cryo-ET) to resolve the first structure of a green algal mitoribosome, stunningly different from both its prokaryotic ancestor, as well as from the flowering plant mitoribosome (Waltz et al., 2020a), but also from all other characterized mitoribosomes across diverse species (Waltz and Giegé, 2020). Our structure reveals how the reduced and fragmented rRNAs are organized and stabilized in the mitoribosome via numerous Chlamydomonas-specific r-proteins. Cryo-ET reveals the native structure and organization of Chlamydomonas mitoribosomes inside mitochondria, revealing that these mitoribosomes are exclusively bound to the inner membrane of mitochondria. Our study provides the first example of a ribosome composed of numerous rRNA fragments, revealing a strikingly divergent blueprint for building this conserved molecular machine.

## Results

### Isolation, mass spectrometry, and cryo-EM of mitoribosomes

To analyze the *C. reinhardtii* mitoribosome, mitochondria were purified and used for mitoribosome isolation following a procedure based on sucrose density gradient separation (see Methods) (Fig. S1). Collected fractions were systematically analyzed by nano-LC MS/MS (Table S1) and screened by cryo-EM to determine their composition. This approach allowed us to identify fractions containing the two mitoribosome subunits which were subsequently used for data collection (Fig. S1). The proteomic analysis identified putative Chlamydomonas-specific r-proteins that were then confirmed by the corresponding cryo-EM reconstructions. Following image processing and extensive particle sorting, reconstructions of both dissociated subunits were obtained. The large subunit (LSU) was resolved to 2.9 Å while the small subunit (SSU) was reconstructed at 5.49 Å and further refined to 4.19 Å for the body and 4.47 Å for the head using a focused refinement approach (Fig. S2). Fully assembled mitoribosomes were identified by nano-LC MS/MS in the cytoribosome fraction (Fig. S1), but cryo-EM investigation revealed aggregates in this fraction, most likely corresponding to mitoribosomes. Nevertheless, the subunit reconstructions were docked into the map of the entire *C. reinhardtti* mitoribosome obtained from the subtomogram averaged mitoribosomes of the *in situ* cryo-ET data (see later in (Fig. 6)) allowing accurate positioning of both subunits relative to each other in the context of a fully assembled native mitoribosome. The isolated subunit reconstructions were similar to the *in situ* subtomogram average, demonstrating that they represent the mature LSU and SSU and not assembly intermediates. Notably, all densities corresponding to novel r-proteins were present in both single particle and subtomogram average reconstructions.

### Overall structure of the Chlamydomonas mitoribosome

Our cryo-EM reconstructions, along with our extensive MS/MS analyses, allowed us to build atomic models of both *C. reinhardtii* mitoribosome subunits (see Methods) (Fig. 1 & 2). The overall architecture of this mitoribosome (Fig. 1) is clearly distinct from both its bacterial ancestor and the flowering plant mitoribosome (Waltz et al., 2020a). Chlamydomonas-specific proteins and domains largely reshape both subunits. Similar to all previously described mitoribosomes, the Chlamydomonas mitoribosome has more r-proteins compared to its bacterial counterpart (Kummer and Ban, 2021; Waltz and Giegé, 2020). These proteins include bacteria-conserved (ancestral r-proteins), r-proteins shared with other mitoribosomes (mitoribosome-specific) and Chlamydomonas-specific r-proteins. In total, the mitoribosome contains 47 r-proteins in the LSU and 36 in the SSU, more than 80 proteins altogether compared to the 54 r-proteins in bacterial ribosomes (Fig. 2 and Table S1). As a result, very few rRNAs are exposed to the solvent, with proteins coating the entire mitoribosome and stabilizing the fragmented rRNAs (Movie 1). Proteins follow the classical r-protein nomenclature (Ban et al., 2014), and newly identified proteins are numbered according to the last inventory of mitoribosomal r-proteins (Valach et al., 2021).

**Figure 1.**
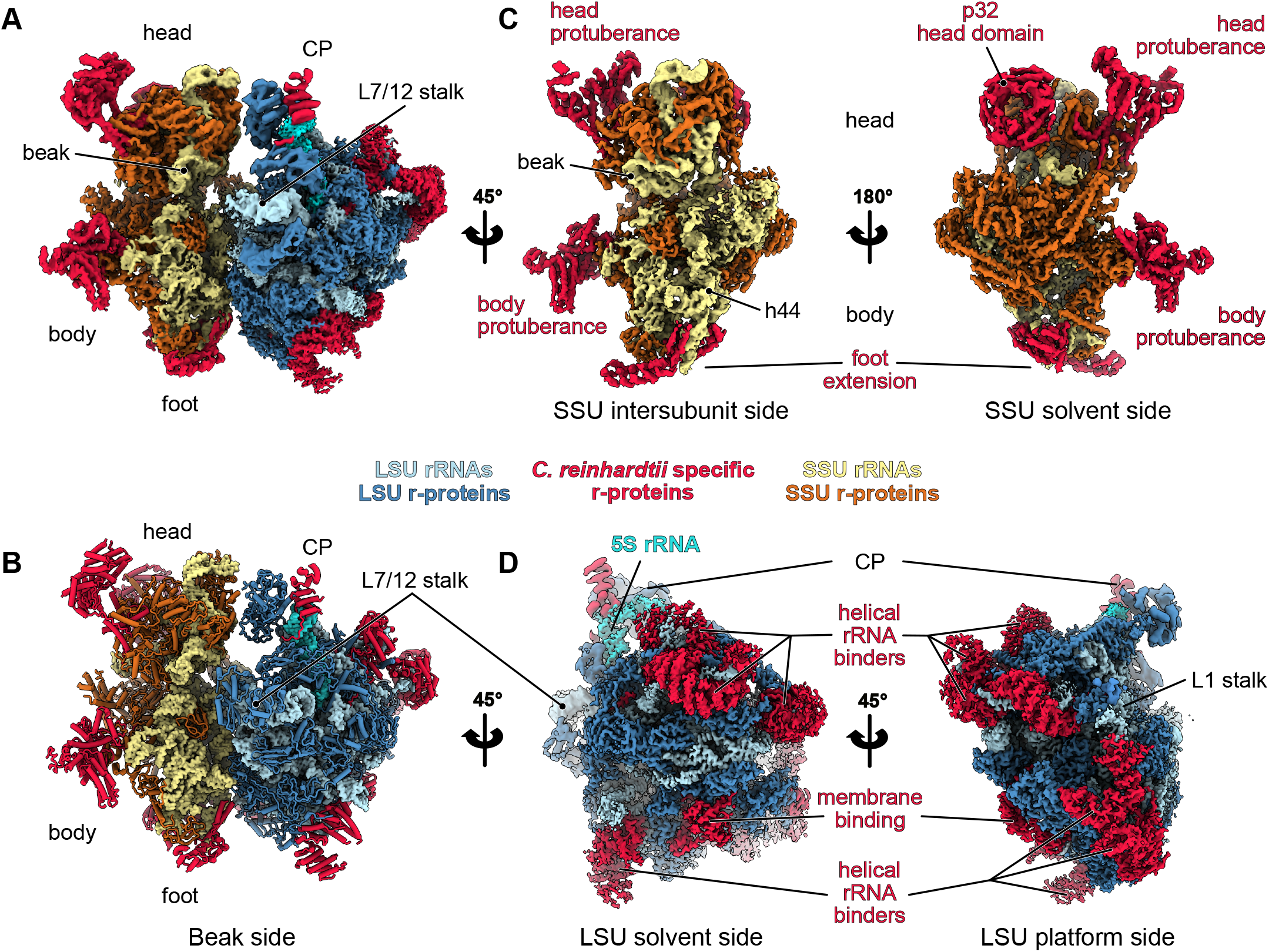
Overall structure of the Chlamydomonas mitochondrial ribosome. **A)** Composite cryo-EM map of the *Chlamydomonas reinhardtii* mitochondrial ribosome and **B)** the resulting atomic model. The large subunit (LSU) components are depicted in blue shades, the small subunit (SSU) components in yellow shades, and the specific r-proteins and domains are displayed in red. **C-D)** Different views of the cryo-EM reconstructions of the SSU (**C**) and the LSU (**D**).

**Figure 2.**
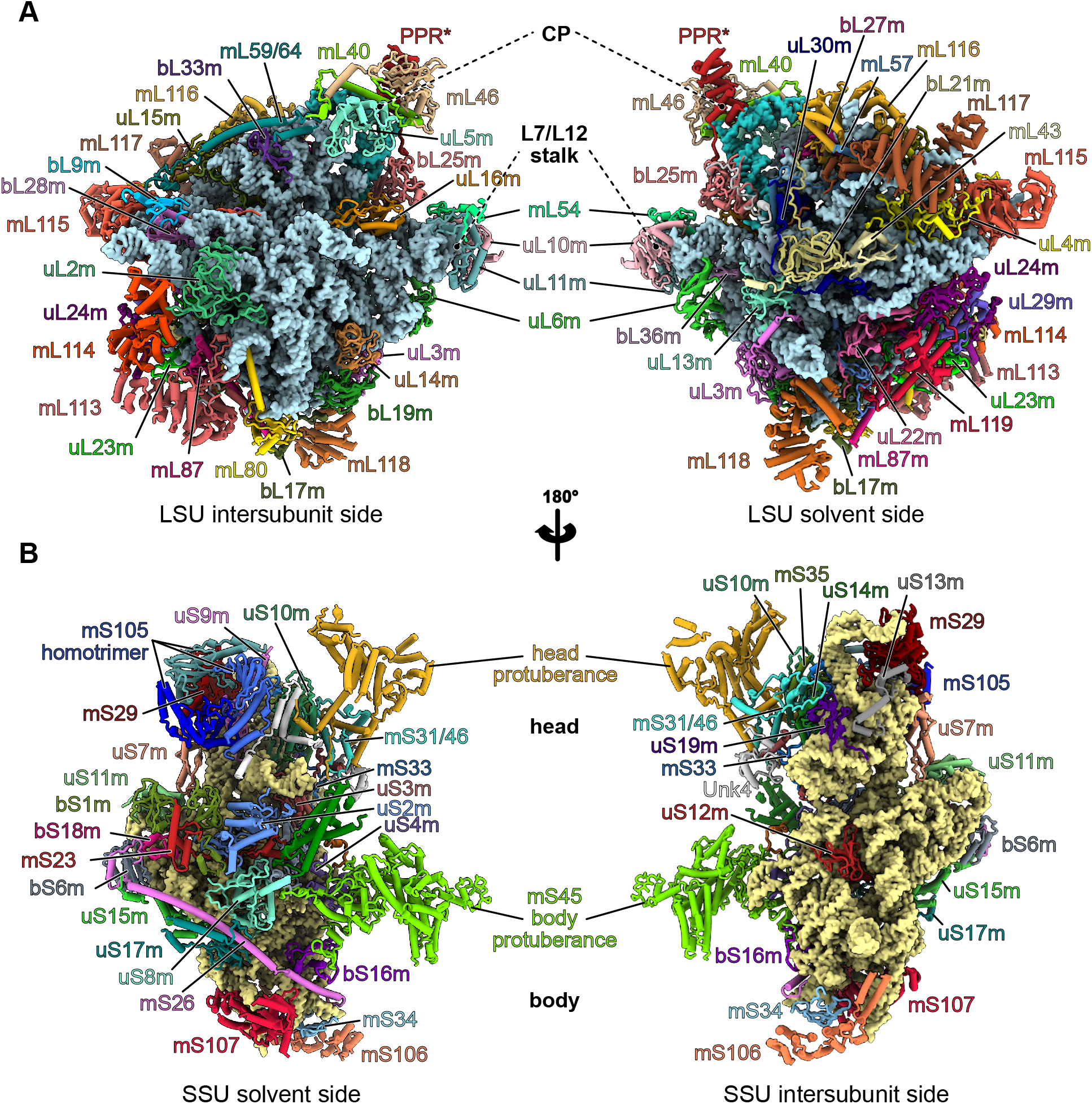
Ribosomal proteins of the Chlamydomonas mitoribosome. **A-B)** The atomic models of the Chlamydomonas mitoribosome LSU (**A**) and SSU (**B**), with mitoribosomal proteins shown in cartoon representation and individually colored and annotated. rRNAs are shown in surface representation and colored in light blue (LSU) and beige (SSU).

Reconstruction of the LSU (Fig. 1D) revealed eight novel proteins (Fig. 1, 2 and Fig. S3 and Table S1). These newly identified proteins are distributed across the whole LSU, where they extend into the solvent and are anchored to the ribosome by interacting with both conserved r-proteins and rRNA fragments. With the exception of mL119 at the exit of the peptide channel, all these proteins are relatively large RNA binders composed of repeated alpha-helical folds, including a mitochondrial TERmination Factor (mTERF) protein, several OctotricoPeptide Repeat (OPR) proteins and one putative PentatricoPeptide Repeat (PPR) protein, named PPR*.

The small subunit (Fig. 1C) reconstruction highlights several distinctive features. Most strikingly, the SSU is shaped by two large protuberances positioned on its beak side, one on the head and one on the body, both formed by helical-rich proteins. The body protuberance, located close to the mRNA entrance, is mainly formed by a specific extension of more than 400 aa in the mitochondria-specific r-protein mS45 (Fig. S4), a highly variable r-protein (Valach et al., 2021). The head protuberance density could not be assigned due to the low resolution of this area. However, several conserved proteins of the SSU head located nearby, notably uS3m, uS10m and mS35, present large extensions. Therefore, it is likely that these extensions could come together and form the head protuberance, as no other apparent candidates could be identified by MS/MS analyses. On the solvent side of the head, a large torus-shaped domain protrudes in the solvent. This additional domain is a homotrimeric complex formed by three copies of mS105, also called p32 (Fig. S4A). This MAM33-family protein is seemingly conserved in all eukaryotes, and has been described to have several functions in mitochondria, some related to mitoribosome assembly (Hillman and Henry, 2019; Jiang et al., 1999; Summer et al., 2020; Yagi et al., 2012). However, this is the first time that mS105/p32 has been identified as a core component of a ribosome. Additionally, the head of the SSU is also characterized by its missing beak, which is typically formed by helix 33 at the junction site of rRNA fragments S3 and S4 (Fig. 3C and Fig. S6). The foot of the SSU is reshaped by Chlamydomonas-specific r-proteins. The extension is formed by two super-helical proteins, one PPR (mS106) and one OPR (mS107). The mS106 protein occupies a position similar to mS27 in humans (Amunts et al., 2015; Greber et al., 2015) and fungi (Itoh et al., 2020), but it does not appear to interact with RNA, nor does it share any sequence identity with mS27 (Fig. S4D). On the other hand, the OPR mS107 directly interacts with rRNA fragment S2, where it encapsulates the tip of helix 11 (Fig. S4C).

**Figure 3.**
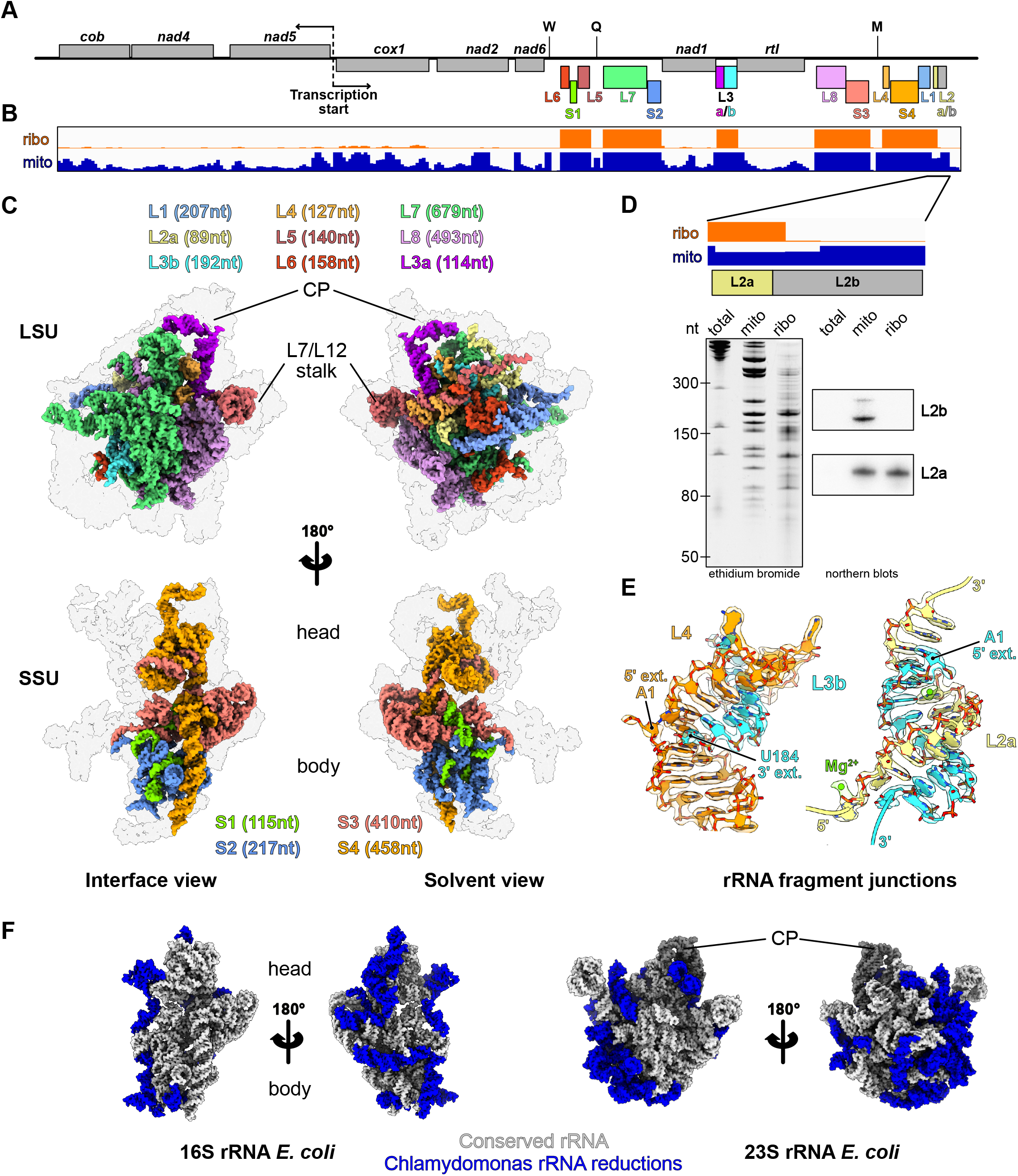
The ribosomal RNAs of the Chlamydomonas mitoribosome are fragmented. **A)** Schematic representation of the entire *C. reinhardtii* mitochondrial genome. The protein-coding genes are displayed in grey and the rRNA fragments incorporated in the mitoribosome are individually colored. The tRNA genes are indicated by letters. **B)** Browser view of the RNAseq data of libraries built from purified mitochondria (orange) and purified mitoribosome (blue) fractions. Data range is [0-700], mapped to the Chlamydomonas mitochondrial genome in (**A**). **C)** rRNA fragment 3D organization in the LSU and the SSU. The r-proteins are shown as a grey silhouette. The fragment colors match the color code used in (**A**). **D)** Zoomed view of the L2a/b coverage (top). The L2b fragment is absent in the ribosomal fraction in the RNAseq analysis. This is confirmed by northern blots (bottom) performed for the L2a and L2b fragments on total cell extracts, mitochondrial and mitoribosomal fractions, and the cryo-EM reconstruction. **E)** Detailed view of the rRNA extremities of the L3b fragment and its pairing with the L2a and L4 fragments. The atomic models are shown mapped into the cryo-EM densities. **F)** Comparison of the Chlamydomonas mitoribosome rRNAs with the *E. coli* ribosome. rRNA reductions are shown in blue.

### Fragmented ribosomal RNAs are assembled to reconstitute the core of Chlamydomonas mitoribosome

In contrast to flowering plants, where rRNAs are largely expanded, the *C. reinhardtii* mitoribosome is characterized by its reduced and fragmented rRNAs (Fig. 3). These rRNAs are scrambled in the mitochondrial genome (Fig. 3A), where they are expressed as a single polycistron which is then further processed into matured transcripts by currently unknown endonucleases (Boer and Gray, 1988). The “23S” and “16S” rRNAs are respectively split into eight fragments totaling 2035 nt, and four fragments totaling 1200 nt (Fig. 3C and Fig. S5 and S6 and Movie 1). This corresponds to 30 % and 22 % reductions compared to bacteria (Fig. 3F). Among all the rRNA pieces predicted to be integrated into the mature mitoribosome, all but one (L2b) could be identified in our cryo-EM reconstructions. To confirm the absence of the L2b fragment, its occurrence was analyzed by comparative RNAseq analyses of mitochondrial and mitoribosomal fractions (Fig. 3B). Consistently, all rRNA fragments could be identified in the mitoribosome fraction except L2b. However, this analysis confirmed that the fragment is indeed expressed and accumulates in the purified mitochondria fraction (Fig. 3B), which is in line with previous transcriptomic analyses (Gallaher et al., 2018; Salinas-Giegé et al., 2017; Wobbe and Nixon, 2013). Additionally, these results were confirmed by RNA blots hybridized against L2b, and an L2a control found in the ribosome and stabilized by the r-protein mL116 (Fig. 3D). Therefore, the L2b RNA is not associated with the mitoribosome, suggesting that this small RNA has an independent function that remains to be elucidated.

In the SSU, the fragmented rRNAs form only a few interactions with the additional r-proteins and are mainly stabilized by base pairing with each other (with the exception the S2 fragment’s h11, which is encapsulated by mS107 (Fig. S4 and S6)). Fragments S1, S2 and a small portion of S3 form the 5’ domain, with the rest of S3 making up domain C. The 3’ end of S3 and the entirety of S4 constitute domains 3’M and 3’m, with S4 largely contributing to linking the head and body of the SSU (Fig. 3).

In contrast to the SSU, the nine rRNA fragments in the LSU are all stabilized by the newly identified Chlamydomonas-specific r-proteins. These fragments reconstitute the different domains of the large subunit. L1 forms the highly reduced domain I of the LSU. L2a, L3b, L4, L5 and part of L6 together form domain II. Portions of L6 and L7 form the highly reduced – almost deleted – domain III. Fragments L7 and L8, the largest of all, make up domains IV, V and VI which form the catalytic core of the ribosome. These three domains are the least altered, with only few helices missing and two expansion segments ES-66 and ES-94 (Fig. 4, Fig. S5), which is most likely due to the high selective pressure to conserve the catalytic region of the ribosome (Petrov et al., 2019). These rRNA fragments are held together by base-pairing with each other, and their extremities are stabilized by base-pairing with other fragments, e.g. L2a, L3b and L4 (Fig. 3E) or with themselves. However, several single stranded rRNA extremities are also stabilized by the Chlamydomonas-specific r-proteins (see below). Surprisingly, while it was anticipated that 5S rRNA should be absent from Chlamydomonas mitoribosome, we identified an RNA density at the typical position of the 5S rRNA in the central protuberance (CP) (Fig. 1 and Fig. S7). This rRNA density could be attributed to the L3a rRNA fragment (Fig. S7). Its association with the mitoribosome was also confirmed by comparative RNAseq analysis of mitochondrial and mitoribosomal fractions (Fig. 3B). Previous studies of the *C. reinhardtii* mitochondrial genome and rRNAs always failed to identify this rRNA as a putative 5S (Boer and Gray, 1988; Denovan-Wright and Lee, 1995; Nedelcu, 1997; Salinas-Giegé et al., 2017). While L3a likely derives from an ancestral bacterial 5S rRNA, it is highly degenerated; very little of the primary sequence is conserved with other 5S rRNAs (Fig. S7D), but a consensus of 6 consecutive nucleotides confirmed its origin. Its overall structure is also weakly conserved, with only the domain γ retaining its characteristic structure to interact with H38. The domain β is angled differently relative to the domain γ stem, which allows the interaction of the terminal loop of the domain β with H87, in contrast to other known ribosome structures. Additionally, the domain α could not be fully resolved, but most likely interacts with the putative PPR protein (labelled “PPR*”), possibly stabilizing its 3’ and 5’ termini. Compared to flowering plants (Waltz et al., 2020a), the Chlamydomonas CP is similar in structure but has retained a canonical bacterial 5S rRNA. However, in terms of composition, the Chlamydomonas mitoribosome CP lacks the uL18m protein, and the specific PPR* protein is present.

**Figure 4.**
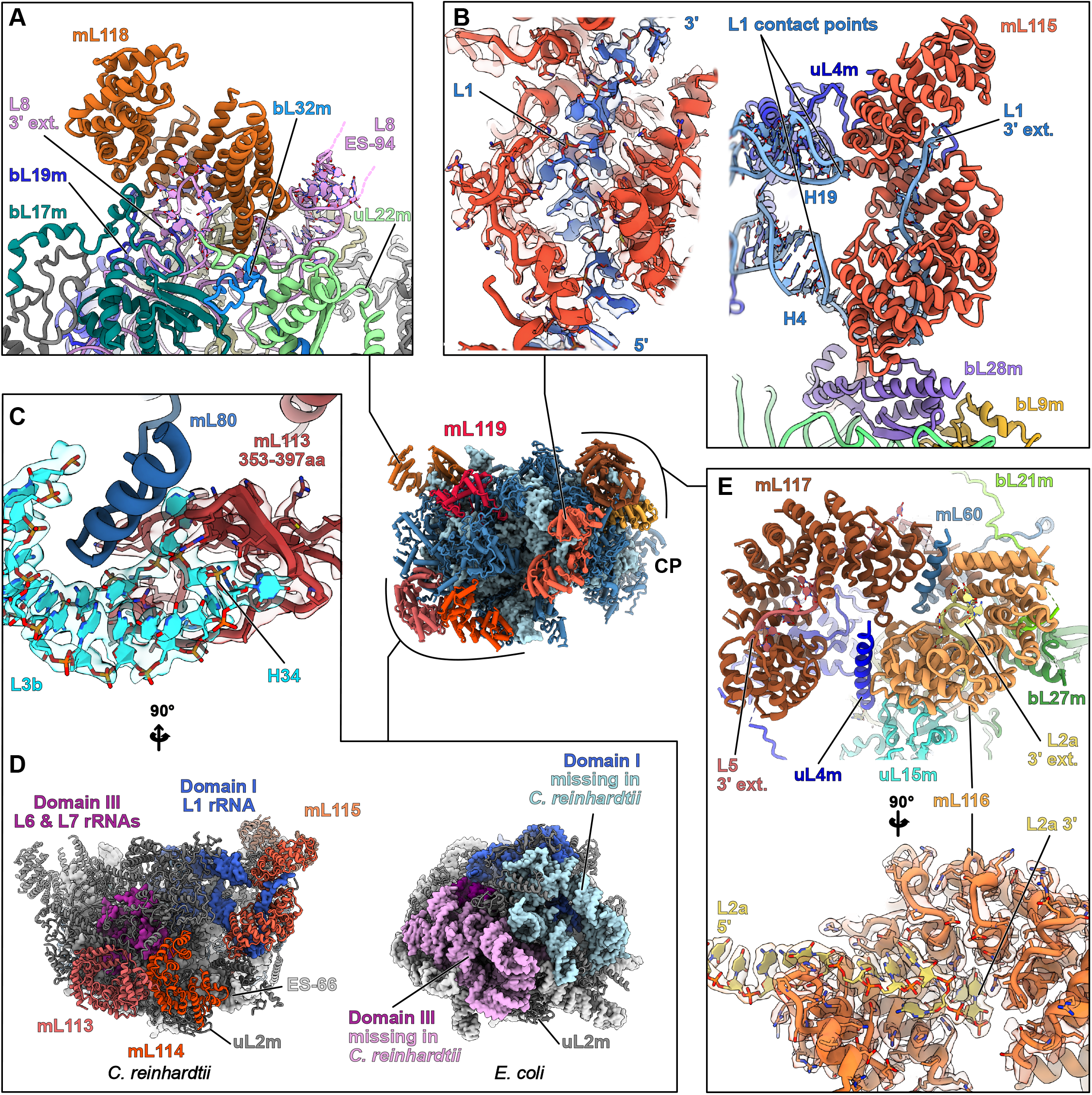
Chlamydomonas-specific proteins stabilize the fragmented rRNAs *via* highly intertwined interactions. Magnified views of the Chlamydomonas-specific r-proteins involved in rRNA stabilization. **A)** mL118 stabilizes the 3’ end of the L8 fragment and contacts the expansion segment 94 (L8 ES-94). **B)** mL115 stabilizes the highly reduced domain I, which is formed by the L1 fragment. mL115 binds the 3’ end of the L1 fragment and also contacts the L1 fragment at two specific points corresponding to H4 and H19. The single stranded portion of L1 interacting with mL115 is shown in its density. **C)** Detailed view of the mL113 contact with the L3b fragment. An inter-repeat domain of mL113 (red) formed by amino acids 353 to 397 clamps the tip of H34 (cyan). The models are shown in their densities. Contrary to the rest of the r-proteins, this one does not enlace single stranded rRNA. **D)** Structural compensation for the loss of large portions of domain I and III. The missing rRNA regions are depicted on the *E. coli* model in pink and light blue, and the compensating proteins mL113, mL114 and mL115 (red shades) are shown on the Chlamydomonas model. mL113 and mL114 compensate for domain III reduction and mL115 stabilizes and compensates the reduced domain I. **E)** The mL116 and mL117 interact with each other and with several surrounding proteins. mL117 is involved in stabilizing the 3’ end of the L5 rRNA fragment, and mL116 stabilizes the L2a 3’ extremity, but also make additional contacts with rRNAs (see Figure S3). The single stranded portion of L2a in mL116 is shown in its density. With the exception of the mTERF protein mL114, the rest of the proteins belong to the same class of ASA2-like/OPR proteins. Further detailed views are shown in Figure S3.

### Specific r-proteins stabilize the rRNAs by highly intertwined protein-RNA interactions

In the LSU, with the exception of mL119, all the Chlamydomonas-specific r-proteins belong to families of predicted nucleic-acid binders. mL114 is an mTERF protein, but mL113, mL115, mL116, mL117 and mL118 all appear to belong to the same protein family as they all have similar tertiary structures and resemble ASA2/OPR proteins. These OPR proteins (OctotricoPeptide Repeat), are predicted to fold into repeated pairs of α-helices, forming a super-helical solenoid, similar to PPR (PentatricoPeptide Repeat) and TPR (TetratricoPeptide Repeat) proteins (Barkan and Small, 2014; Hammani et al., 2014). Both PPR and TPR are widespread in eukaryotes and have previously been found in mitoribosomes, notably in the flowering plants mitoribosome which includes 8 PPR proteins (rPPR proteins), that stabilize the numerous rRNA expansions (Waltz et al., 2020a). In Chlamydomonas, OPR proteins were previously described to be involved in gene expression regulation, notably in the chloroplast (Boulouis et al., 2015; Eberhard et al., 2011; Hammani et al., 2014; Rahire et al., 2012; Wang et al., 2015). Here, these proteins stabilize the many rRNA fragments by different modes of RNA interaction. This is, to our knowledge, the first structural description of this kind of protein in interaction with RNA. Proteins mL115, mL116 and mL117 stabilize the 3’ extremities of L1, L2a and L5, respectively, by binding the single stranded rRNA fragments in their inner groove (Fig. 4). The stabilization is primarily mediated by positive/negative charge interactions, where the inner grooves of the proteins are largely positively charged, filled with lysine and arginine that interact with the negatively charged phosphate backbone of the RNA (Fig. S3). Unlike the rest of the RNA binders, mL113 does not interact with RNA in its inner groove. An inter-repeat domain formed by amino acids 353-397 clamps the tip of H34 from the L3b fragment (Fig. 4C and Fig. S3A). Here, the interaction does not stabilize the rRNA itself, but rather constitutes an anchor point between mL113 and the ribosome. mL118 acts similarly to the SSU’s OPR mS107, as it binds the 3’ extremity of the L8 fragment, which forms a loop inside the inner groove of the protein (Fig. 4A). Additionally, these proteins also interact with RNA *via* the convex side of their super-helical fold. This is the case for mL115 which interacts with H4 and H19 of the L1 fragment. mL116 interacts with H28 of the L2a fragment and H38 formed by the L3b/L4 duplex, and mL118 interacts with ES-94 of the L8 fragment (Fig. S3). Moreover, the mTERF protein mL114 stabilizes the ES-66 *via* its C-terminal region, not its inner groove (Fig. S3A). All these proteins largely interact with conserved proteins as well as with each other, e.g. mL113 with mL114 and mL116 with mL117 (Fig. 4). Furthermore, mL113, mL114 and mL115 structurally compensate missing rRNA on the backside of the LSU (Fig. 4D). mL113 and mL114 compensate for the almost wholly deleted domain III, while mL115 both stabilizes and compensates for the missing parts of domain I. Interestingly, mL113 and mL114 are similarly positioned compared to mL101 and mL104, with the mitoribosomes’ flowering plants rPPR proteins stabilizing the remodeled domain III (Waltz et al., 2020a).

### Knock-down of Chlamydomonas-specific r-proteins affects fitness and rRNA stability

Next, we used targeted gene silencing to investigate the importance of the Chlamydomonas-specific r-proteins for ribosome integrity. We explored the CLiP Chlamydomonas mutant library (Li et al., 2019), but no mutant strains for the genes of interest could be confirmed. Hence, we generated artificial miRNA (amiRNA) strains for the novel factors. This method reduces targeted protein expression at the transcript level (Molnar et al., 2009). Strains were generated for mL113, mL116, mL117, mL118, and mS105 (p32). The physiological phenotype of each amiRNA strain was analyzed, particularly the capacity to grow under heterotrophic conditions (dark + acetate), which is typically defective in Chlamydomonas mutants impaired in mitochondrial respiration (Salinas et al., 2014). Some transformants revealed growth retardation when cultivated in heterotrophic conditions and the ones presenting the most severe macroscopic phenotypes were selected (Fig. 5A). The expression of targeted mRNAs was monitored by quantitative RT-PCR, revealing down-regulations of 78%, 48%, 84%, 51% and 77% on average, respectively (Fig. 5B). The transformants exhibiting the most severe growth phenotypes were selected (Fig. 5A). The expression of targeted mRNAs was monitored by quantitative RT-PCR, revealing average down-regulation of 78%, 48%, 84%, 51% and 77% for mL113, mL116, mL117, mL118, and mS105, respectively (Fig. 5B).

**Figure 5.**
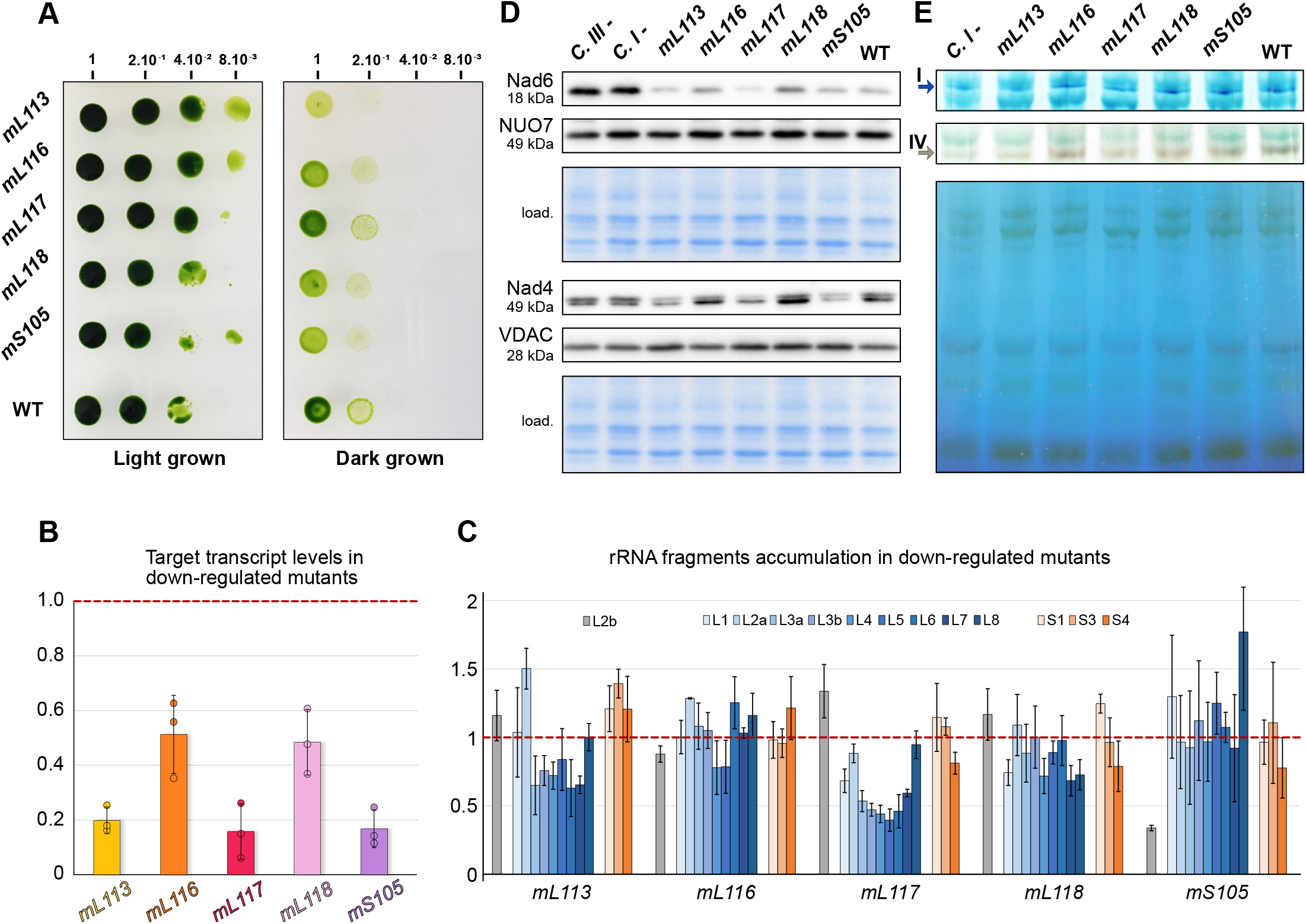
Down regulation of Chlamydomonas-specific r-proteins affect fitness, rRNA accumulation and mitochondrial proteins synthesis. **A)** Growth phenotype of amiRNA knockdown strains. After a first-round selection, transformants were obtained for *mL113*, *mL116*, *mL117*, *mL118* and *mS105*. Growth phenotypes in the dark were investigated by 5-fold dilution series. Dilutions were spotted on two identical TAP plates, one placed in light and the other in darkness. **B)** Relative levels of amiRNA-targeted mRNAs investigated by RT-qPCR. All strains show clear down-regulation compared to WT. **C)** Relative steady-state levels of rRNA fragments in the different Chlamydomonas-specific r-protein amiRNA strains. rRNA fragments of the LSU are shown in blue shades, rRNA fragments of the SSU are shown in orange shades, and L2b (not present in the ribosome) is grey. For (**B**) and (**C**), the WT levels are normalized to 1 on the y-axis, and the average was obtained from three biological replicates, each analyzed in three technical replicates. **D)** Steady-state levels of mitochondria-encoded Nad6 and Nad4, as well as nuclear-encoded NUO7 and the mitochondrial porin VDAC, are shown by protein immunoblots. **E)** Blue native PAGE and in-gel activity assays for complexes I and IV. The resulting bands are indicated by arrows. Wild-type (WT), complex I mutant *dum5* (C I-), and complex III mutant *dum11* (C III-) were used as controls.

The levels of rRNA fragments in the transformants with the highest reduction of the targeted transcript were then determined using quantitative RT-PCR to monitor the effect on rRNAs and mitoribosome stability. This analysis showed that the overall relative levels of LSU rRNAs decreased by 13%, 39% and 14% in the *mL113, mL117, mL118* knockdown strains, while the level of L2b RNA, which does not occur in mitoribosomes, followed a different behavior (Fig. 5C). In contrast, relative rRNA levels were not significantly affected in the *mL116* and *mS105* knockdown strains, with the exception of L2b which is reduced to about 70% in *mS105.* Altogether, it appears that Chlamydomonas-specific r-proteins, in particular mL113, mL117 and mL118, are required for the proper stability of the LSU rRNAs. In addition, the accumulation of mitochondria-encoded proteins was investigated by protein immunoblots. Two mitochondrial-encoded components of respiratory complex I, Nad4 and Nad6, were analyzed, and the nuclear-encoded subunit NUO7 of complex I and the mitochondrial porin VDAC were used as a controls (Fig. 5D). Nad4 levels were decreased in *mS105*, while *mL113* and *mL117* had reduced levels of both Nad4 and Nad6. Finally, the accumulation of assembled respiratory complexes, which contain mitochondria-encoded proteins, was investigated by blue native PAGE (BN-PAGE) coupled to in-gel activity assays (Fig. 5E). These tests revealed that the *mL113* strain is impaired in the activity of complexes I and IV, whereas the *mL117* strain also appears to be affected but to a lesser extent, which correlates with the immunoblot assays. Collectively, these analyses show different impacts on the knockdown strains, suggesting non-redundant functions for these r-proteins.

### The Chlamydomonas mitoribosome is tethered to the inner mitochondrial membrane via two protein contact sites

The LSU reconstruction revealed the presence of a specific r-protein, mL119, precisely located at the exit of the peptide channel (Fig. 6). There, this protein forms several contacts with r-proteins uL22m, uL24m, uL29m and bL32m, and with nucleotides 114-124 of the L6 fragment *via* its C-terminal part, which anchors the protein to the ribosome (Fig. 6E). The N-terminal part of mL119, forming most of the protein’s mass, is exposed to the solvent. This protein has no apparent homolog and appears to be restricted to the Chlorophyceae (green alga) lineage. Similar to humans, the Chlamydomonas mitochondrial genome only codes for membrane components of the respiratory chain, except for the *rtl* gene, whose expression and function remain uncertain. In humans and yeast, it was previously shown that mitoribosomes contact the membrane protein insertase Oxa1 *via* r-protein mL45 in human and the linker protein Mba1 in yeast, the two proteins being homologs (Desai et al., 2020; Itoh et al., 2021; Kummer et al., 2018; Ott and Herrmann, 2010; Ott et al., 2006). These two proteins are positioned at the exit of the peptide channel, where they link the ribosome to the membrane by binding Oxa1, allowing direct insertion of nascent proteins into the membrane. Given the position of mL119, one would expect this protein to fulfill a function similar to mL45 and Mba1. To assess the role of mL119 in membrane binding, Chlamydomonas mitoribosomes were directly visualized inside cells using *in situ* cryo-ET (Fig. 6 and Movie 2).

**Figure 6.**
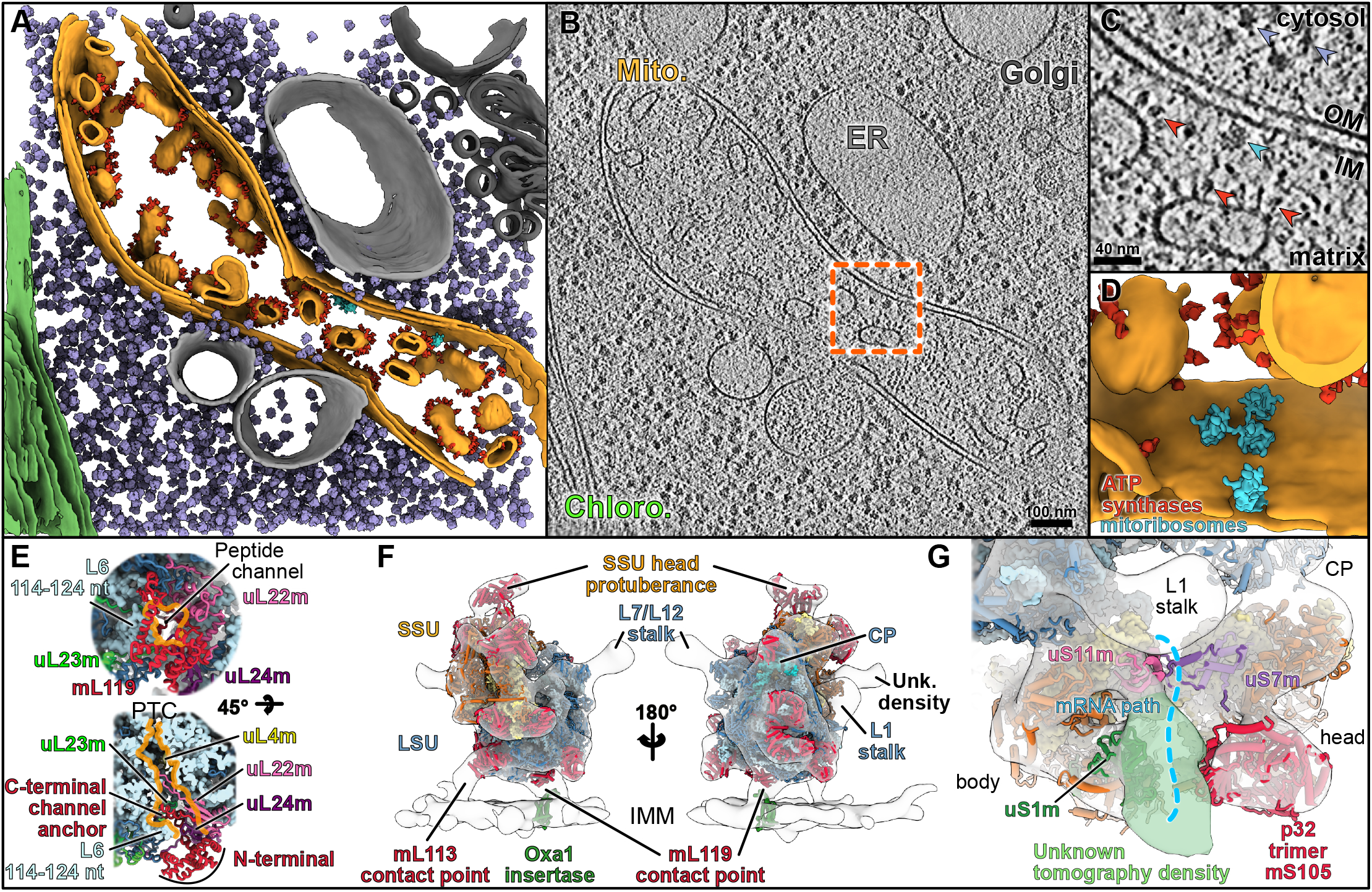
The mitoribosome is attached to the mitochondrial inner membrane *via* specific proteins. **A)** Segmentation of an *in situ* tomogram, depicting a mitochondrion within a native *C. reinhardtii* cell. The mitochondrion is shown in orange, chloroplast in green, ER and lysosome in light gray and Golgi in dark gray. Subtomogram averages of ATP synthase dimers (red), cytosolic ribosomes (purple) and mitoribosomes (cyan) are mapped into the volume. **B)** A slice through the corresponding raw tomogram. **C)** Close-up view of a membrane-bound mitoribosome at the inner membrane (IM), boxed in (**B**). Arrowheads point to ATP synthase, cytosolic ribosomes and a mitoribosome. **D)** Close-up view of the segmented area presented in (**C**), highlighting a cluster of three mitoribosomes that possibly form a polysome. Scale bar are indicated on (**B**) and (**C**). **E)** The peptide channel of the Chlamydomonas LSU from a solvent view (top) and as well as a cut view (bottom). The mL119 protein is located at the exit of the peptide channel. The C-terminal part of mL119 anchors the protein to the ribosome, while the N-terminal part is exposed to the solvent where it could interact with Oxa1. Orange lines delimit the peptide channel. Ribosomal proteins are depicted as cartoons, and rRNAs are depicted in surface representations in light blue. **F)** The atomic model of the mitoribosome is fitted into the subtomogram average density of the native membrane-bound mitoribosome. The mitoribosome contacts the inner mitochondrial membrane (IMM) at two specific points. These contacts are mediated by two Chlamydomonas-specific proteins, mL113 and mL119, the latter of which is located at the exit of the peptide channel, adjacent to the Oxa1 insertase. The model of the bacterial homolog of Oxa1 is shown for illustration purposes. **G)** Close-up view of the mRNA exit channel. The molecular model is fitted into the subtomogram average, highlighting the presence of an additional unknown density (green) located close to uS1m.

Whole *Chlamydomonas reinhardtii* cells were vitrified, thinned by cryo-focused ion beam (FIB) milling, and imaged by cryo-ET. A representative tomogram depicting a section of a native mitochondrion within a *C. reinhardtii* cell is shown in Fig. 6A-D. ATP synthase dimers, cytosolic ribosomes and mitoribosomes were automatically localized by template matching, structurally resolved by subtomogram averaging, and then mapped back into the native cellular environment (Fig. 6B). Contrary to cytosolic ribosomes, which crowd the cytoplasm, mitoribosomes have very low abundance and are localized to the inner mitochondrial membrane. Their low copy number and membrane association highlight the difficulty of purifying these complexes compared to cytosolic ribosomes. Alignment of subtomograms containing mitoribosomes yielded a structure of the native membrane-bound mitoribosome at about 32 Å resolution. With the exception of dynamic flexible regions (e.g. L7/L12 and L1 stalks), the *in situ* subtomogram average is highly similar to our single-particle reconstructions, as revealed by molecular fitting. This structural agreement confirms that the single-particle reconstructions very likely correspond to mature forms of the mitoribosome subunits (Fig. 6F). The *in situ* subtomogram average had one additional density located at the mitoribosomes’s mRNA exit channel. Although we do not know the identity of this density, we speculate that it may correspond to exiting mRNAs or the recruitment of additional factors during active translation (Fig. 6G).

The *in situ* subtomogram average reveals how the mitoribosome is tethered to the inner mitochondrial membrane. Membrane-bound mitoribosomes were previously described by cryo-ET of mitochondria isolated from yeast (Pfeffer et al., 2015) and humans (Englmeier et al., 2017). Similar to yeast, but not humans, the Chlamydomonas mitoribosome makes two distinct contacts with the membrane. Superposition with the atomic model reveals that one contact is located at the precise position of mL119, supporting the hypothesis that this protein could directly interact with the ribosome binding domain of Oxa1 *in vivo*. Therefore, it appears that mL119 constitutes a functional analog of mL45 and Mba1. However, mL119 and mL45/Mba1 are not evolutionary related, but rather appear to have convergently evolution to fulfil the same function. The mitoribosome’s second membrane contact is mediated via the C-terminal part of mL113 (Fig. 6F). This region of mL113 had poorly resolved density in our cryo-EM map and was thus not modeled, but we could still observe its position at low resolution (Fig. S3A). The mL113 contact mimics the rRNA expansion segment ES-96 that forms the second contact site with the membrane in yeast (Pfeffer et al., 2015).

## Discussion

Our study describes the structure and composition of the Chlamydomonas mitochondrial ribosome. The cryo-EM reconstructions show that this green alga mitoribosome differs significantly from prokaryotic ribosomes as well as its flowering plant counterpart (Tomal et al., 2019; Waltz et al., 2020b). In both the small and the large subunits, the mitoribosome has acquired several additional r-proteins that significantly reshape this ribosome’s architecture. These specific r-proteins combined with the fragmented rRNAs (four pieces in the SSU and nine pieces in the LSU) constitute an extreme case of ribosome divergence, even among the exceptionally diverse mitoribosomes.

One striking feature of the SSU is the presence of two large protuberances on the head and the body. The body protuberance could be assigned to a large insertion in the mitoribosome-specific protein mS45 (Fig. S4B), which shows high structural variability between mitoribosomes in different species despite the conservation of its core domain (Valach et al., 2021). The head protuberance was poorly resolved in our density map but is most likely composed of long extensions of the head’s conserved r-proteins (uS3m, uS10m, mS35). In vascular plant mitoribosomes, the uS3m r-protein was shown to form a similar large protuberance on the SSU head, suggesting a common origin of these protrusions (Waltz et al., 2020a). The roles of these two protuberances is unknown. The position of the body protuberance, close to the mRNA entry channel, might suggest a species-specific mechanism of mRNA recruitment, similar to that in humans mediated by mS39 (Aibara et al., 2020; Englmeier et al., 2017; Kummer and Ban, 2021; Kummer et al., 2018). This protuberance, in conjunction with the additional density observed in the subtomogram average next to bS1m at the mRNA exit channel (Fig. 6G), might point to the existence of specific translation processes in Chlamydomonas. Translation initiation in Chlamydomonas mitochondria shares some features with that of human mitochondria, as the mRNAs lack 5’ untranslated regions in both organisms. However, Chlamydomonas most probably has a specific mechanism for translation initiation, as its mitochondrial mRNAs do not have the U-rich motif downstream of the AUG that was proposed to interact with mS39 in humans. Furthermore, mature Chlamydomonas mRNAs have poly-C rich 3’ tails that might be required for translation initiation (Salinas-Giegé et al., 2017).

Another key feature of the SSU is the homotrimeric mS105 (p32) protein forming a torus-shaped protuberance on the back of the SSU head, reminiscent of RACK1 on the cytosolic ribosome (Johnson et al., 2019). This protein belongs to the MAM33 family, which appears to be eukaryote-specific and is characterized by its quaternary structure: a doughnut-shaped trimer that is highly negatively charged (Jiang et al., 1999; Seytter et al., 1998). The p32 protein has been the subject of many studies, as its mutations result in severe diseases in humans (Fogal et al., 2010; Hu et al., 2013; Saha and Datta, 2018; Yagi et al., 2012). However, despite decades of research, its precise functions remain elusive. Recent studies suggest that MAM33-family proteins might be involved in mitoribosome biogenesis. Indeed, they are linked to the LSU biogenesis in yeast (Hillman and Henry, 2019), in Trypanosoma, the recent structure of the SSU “assemblosome” includes a heterotrimeric p22, directly highlighting its role in mitoribosome biogenesis (Saurer et al., 2019; Sprehe et al., 2010) and in humans, the protein YBEY forms a complex with p32 and is involved in the SSU biogenesis (Summer et al., 2020). We observed that p32 is an integral component of the Chlamydomonas mitoribosome. However, it does not appear to have an obvious function related to the translation process, as it does not bind rRNA and only makes a few contacts with the adjacent r-proteins. Likewise, its downregulation does not impair the accumulation of mitochondrial rRNAs. Taking these result together with the above-mentioned studies, it seem that p32 could act in mitoribosome maturation. Given its overall negative charge, it might serve as a binding platform that scaffolds other factors during ribosome biogenesis. It is unclear why p32 would be kept as a constitutive ribosomal protein in Chlamydomonas and not in other eukaryotes, but this may point to species-specific functions.

One of the most prominent features of Chlamydomonas mitoribosome is its fragmented rRNAs, with four pieces in the SSU and nine pieces in the LSU. Kinetoplastida and Euglenozoa also have fragmented rRNA in their cytosolic ribosomes (Greenwood and Gray, 1998; Hashem et al., 2013; Matzov et al., 2020) However, only the LSU rRNA is fragmented, and the fragments are continuous in the genome. It has been proposed that the fragmentation and scrambling of the Chlamydomonas rRNA genes are the result of several mitochondrial genome recombination events between short repeated sequences (Nedelcu, 1997). We assigned all the previously identified rRNA fragments in the mitoribosome except one, L2b. This fragment is not incorporated into the mature mitoribosome and is thus not an rRNA. Nevertheless, its transcript has reproducibly been found in the total mitochondrial fraction (Salinas-Giegé et al., 2017; Wobbe and Nixon, 2013). Its function is unclear, but it might be involved in mitochondrial genome maintenance, which involves telomere-like structures in Chlamydomonas, as the L2b sequence is highly similar to both ends of the linear Chlamydomonas mitochondrial genome (Smith and Craig, 2021; Smith et al., 2010). Importantly, we reveal that the L3a rRNA fragment is a degenerated 5S rRNA, which escaped identification because of its highly divergent primary sequence. Even compared to closely related Chlorophytes and Chlorophyceae species, Chlamydomonas L3a is particularly different at the sequence level (Fig. S7D). L3a occupies the same position as a classical 5S, but its overall structure, notably the domain α and β structures, are quite different from other known 5S structures, rendering it one of the most divergent 5S rRNA described to date. The rest of the rRNA fragments form the core of the mitoribosome and are globally conserved, yet reduced. In the ribosome core, these fragments are stabilized by base-pairing with each other. In contrast, on the outer shell of the ribosome, especially in the LSU, the rRNA fragments are stabilized by the Chlamydomonas-specific r-proteins. These proteins are all alpha-helical repeats belonging to nucleic-acid binder families PPR, OPR and mTERF. In our structures, they form highly intertwined interfaces with single- and double-stranded RNA, all involving positive/negative charge interactions. These proteins stabilize the 3’ end of rRNA fragments L1, L2a, L5 and L8 by enlacing their single stranded extremities and also contact and stabilize additional rRNA helices via their convex surfaces.

The function of these proteins was investigated by analyzing down-regulation mutants (Fig. 5). All mutants for r-proteins of the large subunit, except the *mL116* mutant, show a diminution of the abundance of rRNA fragments of the LSU, indicating that these novel r-proteins are important for the stability of the rRNAs, and thus integrity of the mitoribosome. They seem to play a role of chaperones, stabilizing and perhaps contributing to the recognition and recruitment of the different rRNA pieces during assembly. In *mS105,* where the protein does not directly interact with rRNAs, the rRNA levels are almost unaffected. Additionally, in downregulation mutants *mL113* and *mL117* steady-state levels of mitochondria-encoded proteins and active respiratory complexes is decreased, highlighting their importance in translation and impact on mitochondrial metabolism.

Our cryo-ET analysis reveals that Chlamydomonas mitoribosomes are bound to the inner mitochondrial membrane. In animals and most eukaryotes, the mitochondrial genome encodes almost only components of the respiratory chain, which are all membrane-embedded proteins. These proteins are co-translationally inserted into the inner mitochondrial membrane to reduce the probability of inefficient protein aggregation during transport (Ott et al., 2016). To facilitate this process, mitoribosomes are consistently found attached to the inner membrane (Englmeier et al., 2017; Pfeffer et al., 2015). In mammals, the mitoribosome attachment is mediated by a specific r-protein, mL45, located at exit of the peptide channel, which links the ribosome to the main insertase of the inner membrane, Oxa1 (Desai et al., 2020; Itoh et al., 2021). In yeast, where one of the mitochondria-encoded proteins is soluble (Freel et al., 2015), the association is mediated by an mL45 homolog, Mba1, which is not an integral constituent of the mitoribosome, and an expansion segment of H96 directly contacting the membrane (Desai et al., 2017; Ott and Herrmann, 2010; Ott et al., 2006). This is most likely also the case in the fungi *N. crassa* (Itoh et al., 2020). In Chlamydomonas, similarly to mammals, all proteins encoded in the mitochondrial genome are components of the respiratory chain. Therefore, it is not surprising that the Chlamydomonas mitoribosome would have acquired a specific r-protein to tether translation to Oxa1. Interestingly, the membrane interaction in Chlamydomonas is mediated via two contact points. mL119 forms the first contact, but mL113 create a second contact point, directly with the membrane similarly to ES-H96 in yeast. Notably, despite the similar location of the mL113 and ES-H96 contact sites, they are of different molecular nature (protein vs. rRNA) and have been acquired via different evolutionary mechanisms: an expansion of the nuclear genome in case of mL113 vs. the expansion of the mitochondrial gene coding for 23S rRNA in yeast. In light of recent literature suggesting an early expansion of mitoribosomal proteins in eukaryotes (Gray et al., 2020), this raises the question whether the second contact site displays an isolated case of convergent evolution between green algae and yeast, or whether is a more universal feature of mitoribosomes that was either replaced (yeast) or lost (mammalian mitoribosome) throughout evolution. The fact that membrane association is mediated by different proteins in each organism, yet the Oxa1 contact is conserved, indicates that this interface is particularly critical (Ott and Herrmann, 2010). In flowering plant mitochondria, which still encode a large number of soluble proteins, accessory factors might recruit mitoribosomes to the membrane, similar to Mba1 in yeast. However, in Tetrahymena, the mitochondrial genome encodes numerous soluble proteins, but the mitoribosome has still acquired a probable permanent anchor to Oxa1, the r-protein mL105 (Tobiasson and Amunts, 2020). Finally, contrary to mammals where mL45 blocks the peptide channel until the mitoribosome association with the membrane (Itoh et al., 2021; Kummer et al., 2018), or the assembly factor mL71 in kinetoplastids (Soufari et al., 2020), here the peptide channel is not blocked by mL119, which does not suggest an inhibition of translation until the mitoribosome is membrane tethered.

In conclusion, our structural and functional characterization of Chlamydomonas mitoribosome provides a new perspective on mitoribosome evolution and membrane binding. Our study paves the way for future investigations of mitoribosomes in other species, in particular apicomplexa such as Plasmodium and Toxoplasma, where fragmented mitochondrial rRNAs also occur (Feagin et al., 2012; Waltz and Giegé, 2020). Altogether, the structure presented here provides further insights into the evolution of mitoribosomes and the elaboration of independent new strategies to perform and regulate translation. Truly, the mitoribosome is one of nature’s most eclectic playgrounds for evolving diverse strategies to regulate a fundamental cellular process.

## Methods

### *Chlamydomonas reinhardtii* mitochondria and mitoribosome purification

*Chlamydomonas reinhardtii* cell wall-less strain CC-4351 (*cw15–325 arg7–8* mt+) was used for mitochondria purification and transformation. The mitochondrial mutants *dum5* (Cardol et al., 2003) and *dum11* (Dorthu et al., 1992) were used as a control for the phenotypic growth analysis in the dark, kindly provided by Dr. Remacle (University of Liège) and respectively annotated on figures as CI- and CIII-. The strains were grown on Tris-Acetate Phosphate (TAP) solid or liquid medium (Harris et al., 2009), supplemented with 100 μg/ml of arginine when necessary, under continuous white light (50 μE m−2 s−1), or in the dark. The purified mitochondrial fraction was obtained as in (Salinas et al., 2012) by digitonin treatment followed by a discontinuous Percoll gradient.

Mitoribosome purification was conducted as previously (Waltz et al., 2019, 2020a). Purified mitochondria were re-suspended in Lysis buffer (20 mM HEPES-KOH, pH 7.6, 100 mM KCl, 30 mM MgCl_2_, 1 mM DTT, 1.6 % Triton X-100, 0.5% n-DDM, supplemented with proteases inhibitors (C0mplete EDTA-free)) to a 1 mg/ml concentration and incubated for 15 min in 4°C. Lysate was clarified by centrifugation at 25.000 g, 20 min at 4°C. The supernatant was loaded on a 40% sucrose cushion in Monosome buffer (Lysis buffer without Triton X-100 and 0.1% n-DDM) and centrifuged at 235.000 g, 3h, 4°C. The crude ribosomes pellet was re-suspended in Monosome buffer and loaded on a 10-30 % sucrose gradient in the same buffer and run for 16 h at 65,000 g. Fractions corresponding to mitoribosomes were collected, pelleted and re-suspended in Monosome buffer and analyzed by nanoLC-ESI-MS/MS and cryo-EM (Fig. S1).

### Grid preparation

For the single particle analyses, 4 μL of the samples at a protein concentration of 1.5 μg/μl was applied onto Quantifoil R2/2 300-mesh holey carbon grid, coated with thin home-made continuous carbon film and glow-discharged (2.5 mA for 20 sec). The sample was incubated on the grid for 30 sec and then blotted with filter paper for 2 sec in a temperature and humidity controlled Vitrobot Mark IV (T = 4°C, humidity 100%, blot force 5) followed by vitrification in liquid ethane.

### Cryo-electron microscopy data collection

The single particle data collection was performed on a Talos Arctica instrument (Thermofisher Company) at 200 kV using the SerialEM software for automated data acquisition. Data were collected at a nominal underfocus of −0.5 to −2.5 μm at a magnification of 36,000 X yielding a pixel size of 1.13 Å for the SSU and 45,000 X yielding a pixel size of 0.9 Å for the LSU. Micrographs were recorded as movie stack on a K2 direct electron detector (GATAN Company), each movie stack were fractionated into 65 frames for a total exposure of 6.5 sec corresponding to an electron dose of 45 ē/Å2.

### Electron microscopy image processing

Drift and gain correction and dose weighting were performed using MotionCorr2 (Zheng et al., 2017). A dose weighted average image of the whole stack was used to determine the contrast transfer function with the software Gctf (Zhang, 2016). The following process has been achieved using RELION 3.0 (Zivanov et al., 2018). Initial analyses were performed in CryoSPARC (Punjani et al., 2017) to asses sample composition and to generate *ab-initio* cryo-EM map. After reference-free 2D classification, for the LSU 346,994 particles were extracted and used for 3D classification into 6 classes (Fig. S2). *Ab-initio* cryo-EM reconstruction generated in CryoSPARC was low-pass filtered to 30 Å, and used as an initial reference for 3D classification. 2 subclass depicting high-resolution features was selected for refinement with 101,291 particles. After Bayesian polishing, the LSU reconstruction reached 3.00 Å resolution. For the SSU reconstruction a similar workflow was applied. After 2D classification 445,469 particles were extracted and used for 3D classification into 6 classes. 1 subclass depicting high-resolution features was selected for refinement with 40,131 particles. After focus refinement using masks for the head and body of the small subunit, the SSU reconstruction reached a resolution of 4.19 Å for the body and 4.47 Å for the head. Determination of the local resolution of the final density map was performed using ResMap (Kucukelbir et al., 2014).

### Structure building and model refinement

The atomic model of the LSU of *C. reinhardtii* was built into the high-resolution maps using Coot, Phenix and Chimera. Atomic models from *E. coli* (PDB: 5KCR) and *A. thaliana* mitoribosome (PDB: 6XYW) were used as starting points for protein identification and modelisation as well as rRNA modelisation. The online SWISS-MODEL (Waterhouse et al., 2018) service was used to generate initial models for bacterial and mitochondria conserved r-proteins. Models were then rigid body fitted to the density in Chimera (Pettersen et al., 2004) and all subsequent modeling was done in Coot (Emsley et al., 2010). Extensions were built has polyalanine and mutated to the adequate sequences. Chlamydomonas-specific proteins for which no model could be generated were first built entirely as polyalanine, then the sequence-from-map PHENIX tool (Liebschner et al., 2019) was used to identify each of the proteins, and the correct sequences were placed in the densities. For the SSU, due to the lower resolution in comparison to the LSU, all extensions of the homology models were built as polyalanine and unknown densities were built as Ala residues. For refinement, a combination of regularization and real-space refine was performed in Coot for each proteins. The global atomic model was then subjected to real space refinement cycles using *phenix.real_space_refine* PHENIX (Liebschner et al., 2019) function, during which protein secondary structures, Ramachandran and side chain rotamer restraints were applied. Several rounds of refinement (manual in Coot and automated using the *phenix.real_space_refine*) were performed to obtain the final models, which were validated using the built-in validation tool of PHENIX, based on MolProbity. Refinement and validation statistics are summarized in Supplementary Table 1.

### Cell Vitrification and Cryo-FIB Milling

For FIB-milling, *Chlamydomonas reinhardtii* mat3-4cells (strain CC-3994) (Umen and Goodenough, 2001), which exhibit superior vitrification due to their small size, were used. They were acquired from the Chlamydomonas Resource Center, University of Minnesota, St. Paul. Cells were grown until mid-log phase in Tris-acetate-phosphate (TAP) medium under constant light exposure and bubbling with normal atmosphere. Vitrification and FIB sample preparation were performed as previously described (Schaffer et al., 2015, 2017). Using a Vitrobot Mark 4 (FEI), cells in suspension (4 μl of ~1,000 cells per μl) were blotted onto R2/1 carbon-coated 200-mesh copper grids (Quantifoil Micro Tools) and plunge frozen in a liquid ethane/propane mixture. Grids were then mounted into Autogrid supports (FEI) and transferred into either a FEI Scios or FEI Quanta dual-beam FIB/SEM instrument. The grids were coated with an organometallic platinum layer by the gas injection system (FEI), and cells were thinned from both sides with a gallium ion beam to a final thickness of ~100–200 nm.

### Cryo-electron tomography data acquisition

Cellular tomograms were acquired on a 300 kV Titan Krios microscope (FEI), equipped with a Gatan post-column energy filter (968 Quantum) and a direct detector camera (K2 summit, Gatan) operated in movie mode at 12 frames per second. Tilt series were recorded using SerialEM software (Mastronarde, 2005) with 2° tilt increments from −60° to +60° (in two halves separated at either 0° or −20°), an object pixel size of 3.42 Å, a defocus of −4 to −5.5 μm, and a total accumulated dose of <100 e−/Å.

### Cryo-electron tomography data processing

Movies from the K2 detector were motion corrected with MotionCor2 (Zheng et al., 2017). Using IMOD software, tilt series were aligned with patch-tracking, and tomograms were reconstructed with weighted back projection. Out of ~130 tomograms, 47 tomograms containing mitochondria were selected. An initial structure of the *C. reinhardtii* mitoribosome was obtained by manually picking 104 mitoribosomes from 27 tomograms following template-free alignment by spherical harmonics (Chen et al., 2013). The initial map was then used as a template for automated template matching on 47 tomograms with a voxel size of 2.1 nm using PyTOM (Hrabe et al., 2012). To reduce-false positives, the highest correlation peaks of the resulting 6-D cross-correlation function localized in the mitochondrial matrix were manually inspected in UCSF Chimera (Pettersen et al., 2004), and a set of 228 subvolumes from 27 tomograms was obtained. The subvolumes were reconstructed at a voxel size of 6.84 Å, aligned using PyTOM’s real-space refinement, and subjected to 1 round of classification with a mask encompassing the membrane region. This yielded a class of 73 mitoribosomes with a clear membrane-density that was subjected to one more round of real-space refinement in PyTOM. For the resulting average, a resolution of 32 Å (large ribosomal subunit) and 34 Å (small ribosomal subunit) was determined by fourier-shell cross-resolution of the two map against the maps obtained by single particle analysis (FSC = 0.33). For the localization of ATP synthases, an initial structure of the *C. reinhardtii* ATP synthase was obtained by manually picking 417 subvolumes from four tomograms and aligning them using spherical harmonics. The obtained map was used as template for automated template-matching on tomograms with a voxel size of 2.1 nm using Pytom, and the highest correlation-peaks were then manually inspected in Chimera to remove false-positives. Cytosolic 80S ribosomes were localized in one tomogram by filtering EMDB-1780 (Armache et al., 2010) to 40 Å and using it as a template for template matching on deconvolved tomogram (tom_deconv; https://github.com/dtegunov/tom_deconv) using Pytom. Subvolumes were extracted for the 2,100 highest correlation peaks, of which 1,700 subvolumes were classified as ribosomes by unsupervised, autofocused 3D classification (Chen et al., 2014).

### Proteomic analyses of *C. reinhardtii* mitoribosome composition

Mass spectrometry analyses of the total, mitochondrial and ribosomal fractions of *C. reinhardtii* were done at the Strasbourg-Esplanade proteomic platform and performed as previously (Waltz et al., 2019). In brief, proteins were trypsin digested, mass spectrometry analyses and quantitative proteomics were carried out by nanoLC-ESI-MS/MS analysis on a QExactive+ (Thermo) mass spectrometer. Data were searched against the UniProtKB (Swissprot+trEMBL) database restricted to the *C. reinhardtii* taxonomy with a target-decoy strategy (UniProtKB release 2020_03, taxon 3055, 31246 forward protein sequences), Proteins were validated respecting FDR<1% (False Discovery Rate) and quantitative label-free analysis was performed through in-house bioinformatics pipelines.

### Artificial miRNA *C. reinhardtii* strain generation and analyses

Artificial microRNAs constructs were created according to (Molnar et al., 2009). The oligonucleotides were designed using the WMD3 Web MicroRNA Designer software v3.2 (wmd3.weigelworld.org) and genome release Chlamydomonas CDS reinhardtii 281 v5.6.cds (Phytozome) (Table S3). The oligonucleotides were annealed, phosphorylated, and ligated into a *SpeI*-digested pChlamiRNA2 containing the ARG7 gene as a selection marker. The resulting plasmids were linearized and transformed into Chlamydomonas CC-4351 strain by the Neon® Transfection System (LifeTechnologies) according to the GeneArt® MAX Efficiency® Transformation protocol for Algae (LifeTechnologies Cat#A24229). Cells with integrated plasmid were selected on TAP plates without arginine. Colonies (16 to 48 depending on the transformation) were picked to grown to logarithmic phase on liquid TAP medium. They were then spotted on two identical TAP plates to test their capacity to grow in the dark. One plate was placed in a mixotrophic condition (light + acetate) for 5-7 days, and the other one in a heterotrophic condition (dark + acetate), for 10-15 days. For the dilution series, cells were grown for 3-4 days on TAP plates and were resuspended in 2 ml of liquid TAP medium. The cell density was measured spectrophotometrically at OD750 and diluted to an OD750 = 1.5. This normalized suspension was used as the starting material (set to 1) for making three serial 5-fold dilutions (2.10^−1^, 4.10^−2^, and 8.10^−3^). A volume of 10 μl for each dilution was then spotted on two identical TAP plates.

### rRNA analysis by RNA sequencing

The RNAs were prepared from cells using TRI Reagent® (Molecular Research Center) according to the manufacturer’s instructions.

For northern blots, 1 μg of total, mitochondrial and mitoribosome fraction RNA, were separated on 7M Urea - 8% polyacrylamide gel, transferred onto Amersham Hybond™-N+ membrane (GE Healthcare Cat#RPN203B) and hybridized to radiolabelled oligonucleotide probes (Table S3) in 6 x SSC, 0.5% SDS at 45°C. Washing conditions were: 2 times 10 min in 2 x SSC and 1 time 30 min in 2 x SSC, 0.1% SDS at the hybridization temperature. For each specific probe, the signal was detected with the Amersham Typhoon laser scanner (Amersham).

For the quantitative real-time RT-PCR analyses, RNAs were treated with RQ1 RNase-Free DNase (Promega Cat#M6101) according to Promega’s protocol, using 0.2 U/μg of RNA. To obtain cDNA, reverse transcription assays were performed according to the manufacturer’s instructions with 2,5 μg of total RNA in the presence of 5 μM of oligo(dT) primer (Table S3) and 25 ng/μl of Random Primers (Promega Cat#C118A) using the SuperScript™ IV Reverse Transcriptase (Invitrogen Cat#18090010). The RT-qPCR amplification was carried out with the dsDNA-specific dye Takyon™ SYBR® 2X qPCR Mastermix Blue (Eurogentec Cat#UF-FSMT-B0701) and monitored in real-time with a LightCycler 480 instrument (Roche). The primers used are listed in (Table S3). The delta-delta Ct method was used to calculate the relative RNA abundance with respect to the geometric mean of two RNA references *MAA7* and *CYN19-3* (Livak and Schmittgen, 2001).

For the RNA sequencing, the p204 library was built with total mitochondrial RNA. The RNA was first chemically fragmented (4 min) and then enzymatically treated with Antarctic Phosphatase (NEB#M0289S) and T4 Polynucleotide Kinase (NEB#M0201S). Library preparation was done according to the TruSeq Small RNA Sample Preparation Guide #15004197 Rev. F February 2014. The library was sequenced on the Illumina MiSeq sequencer in a paired-end mode of 2X75 nt reads. The NGS192-small library was built with the mitoribosome fraction. The RNA was also enzymatically treated with Antarctic Phosphatase and T4 Polynucleotide Kinase. The library was then constructed with the NEBNext multiplex small RNA Library set for Illumina reference E7580 following the manufacturer’s instructions. Following PCR amplification, a size selection was performed on a 6% TBE gel to recover the 160-350 bp PCR fragments for sequencing. The NGS192-total library was prepared according to the Truseq Stranded Total RNA with Ribozero Plant kit, starting from the first-strand cDNA synthesis step and omitting the two first depletion and fragmentation steps. The library was sequenced on the Illumina MiSeq sequencer in a paired-end mode of 2X110 nt reads. Both libraries were sequenced at the IBMP platform. The reads were mapped to the Chlamydomonas mitochondrial genome (EU306622) using Bowtie2 version 2.4.1 with the following options --end-to-end --very-sensitive -N 0 -L 22. Alignments were displayed with the Integrative Genomics Viewer (IGV) with the bigWig format.

### Protein analyses

The Chlamydomonas crude total membrane fractions were obtained according to (Remacle et al., 2001). Chlamydomonas cells TAP liquid cultures were collected and resuspended to 2-1.6×108 cells/ml in MET buffer (280 mM mannitol, 0.5 mM EDTA, 10 mM Tris-HCl pH 7) with 1x cOmplete™ Protease Inhibitor Cocktail and then disrupted by sonication (four times 30 sec of sonication and 30 sec of pause; Bioruptor® Pico, Diagenode). The suspension was centrifugated 10 min at 500 g, followed by 4 min at 3000 g) and the protein content of the supernatant was determined by the Bradford method. Equal amounts of protein were separated using 15% SDS−polyacrylamide gel electrophoresis (PAGE), and transferred to a 0.45 μm PVDF membrane (Immobilon®-P Transfer Membrane; Merck Millipore Cat.#IPVH00010). Specific antibodies were used in immunoblotting and were detected using chemiluminescence (Clarity Western ECL Substrate, Bio-Rad). We used rabbit sera obtained against Chlamydomonas reinhardtii mitochondrial-encoded sub-units complex I, Nad4 (1:1000) and Nad6 (1:100), nuclear-encoded sub-unit complex I, NUO7 (1:2000), and the nuclear-encoded mitochondrial protein VDACI (1:25000). Blue native polyacrylamide gel electrophoresis (BN-PAGE) analyses were conducted according to (Schägger et al., 1991). The crude total membrane fractions were prepared as above with an additional centrifugation at high speed (27000 *g* for 15 min) and the final pellet was suspended in ACA buffer (375 mM 6-aminohexanoic acid, 25 mM Bis-Tris, pH 7, and, 250 mM EDTA). 0.5 mg of crude total membrane were first solubilized in the presence of 1,5 % (w/v) n-dodecyl-β-D-maltoside and then centrifuged for 40 min 14200 g at 4°C to remove insoluble matters. 0.65 % (w/v) of coomassie serva blue G was then added to the supernatant prior to separation by electrophoresis on a 5% to 12% polyacrylamide gradient BN gel. In-gel detection of Complex I (NADH dehydrogenase) activity was performed using a 100 mM Tris-HCl pH 7.4 buffer containing 200 μM NADH and 0.2 % nitro blue tetrazolium (NBT). In-gel detection of Complex IV (cytochrome *c* oxydase) activity was performed using a 10 mM MOPS-KOH pH 7.4 buffer containing 7.5 % saccharose, 19 U/ml catalase from bovine liver, 0.1 % cytochrome *c* and 0.01 % 3,3’-diaminobenzidine (DAB).

### Figure preparation and data visualization

For the tomography reconstruction, segmentation of ER, mitochondrial and chloroplast membranes was done using EMAN’s convolutional neural network for automated annotation (Chen et al., 2017). Using the TOM toolbox in matlab (Nickell et al., 2005), the averages of the cytoribosomes, mitoribosomes and ATP synthases were pasted into the tomogram at the refined coordinates and angles determined by subtomogram analysis. Figures featuring cryo-EM densities as well as atomic models were visualized with UCSF ChimeraX (Goddard et al., 2018) and Chimera (Pettersen et al., 2004).

## Data Availability

The cryo-EM maps of *C. reinhardtii* mitoribosome have been deposited at the Electron Microscopy Data Bank (EMDB): EMD-XXXX for the LSU, EMD-XXXX for the head of the SSU, EMD-XXXX for the body of the SSU and EMD-XXXX for the subtomogram averaging of the whole ribosome. The corresponding atomic models have been deposited in the Protein Data Bank (PDB) under the accession XXXX for the LSU and XXXX for the SSU. Mass spectrometric data have been deposited to the ProteomeXchange Consortium via the PRIDE partner repository with the dataset identifier PXD024708 and 10.6019/PXD024708. RNAseq data were deposited in the NCBI Gene Expression Omnibus under accession number GSE171125.

## Acknowledgments

This work has benefitted from the facilities and expertise of the Biophysical and Structural Chemistry platform (BPCS) at IECB, CNRS UMS3033, Inserm US001, University of Bordeaux. We thank, J. Chicher and P. Hamman of the Strasbourg Esplanade proteomic analysis for the proteomic analysis. The mass spectrometry instrumentation was funded by the University of Strasbourg, IdEx “Equipement mi-lourd” 2015. We thank Miroslava Schaffer for help with FIB milling and cryo-ET acquisition. We thank Jürgen Plitzko and Wolfgang Baumeister for access to cryo-EM instrumentation and support. We are grateful to Pr. Claire Remacle (University of Liège) for her kind gifts of strains and antibodies. This work was supported by a European Research Council Starting Grant (TransTryp ID:759120) to YH, by the LabEx consortium ‘MitoCross’ in the framework of the French National Program ‘Investissement d’Avenir’ (ANR-11-LABX-0057_MITOCROSS) to PG and a Agence Nationale de la Recherche (ANR) grant [MITRA, ANR-16-CE11-0024-02] and [DAMIA, ANR-20-CE11-0021] to YH and PG. Additional funding was provided by the Max Planck Society and the Helmholtz Zentrum München.

## Author Contributions

FW, LD, PG and YH designed and coordinated the experiments. HM and TSG generated the amiRNA strains and analyzed them. TSG and FW purified mitochondria. FW purified the mitochondrial ribosomes. HS and FW acquired the cryo-EM data and performed the single particle analysis. RE, SP, FF and BE collected and analyzed the cryo-ET data, including subtomogram averaging. FW built the atomic models and interpreted the structures. LK performed the mass-spectrometry experiments. FW prepared and assembled the figures. FW wrote the manuscript, which was edited by TSG, RE, HS, HM, LK, SP, FF, BE, PG, LD and YH.

---

**Figure S1.**
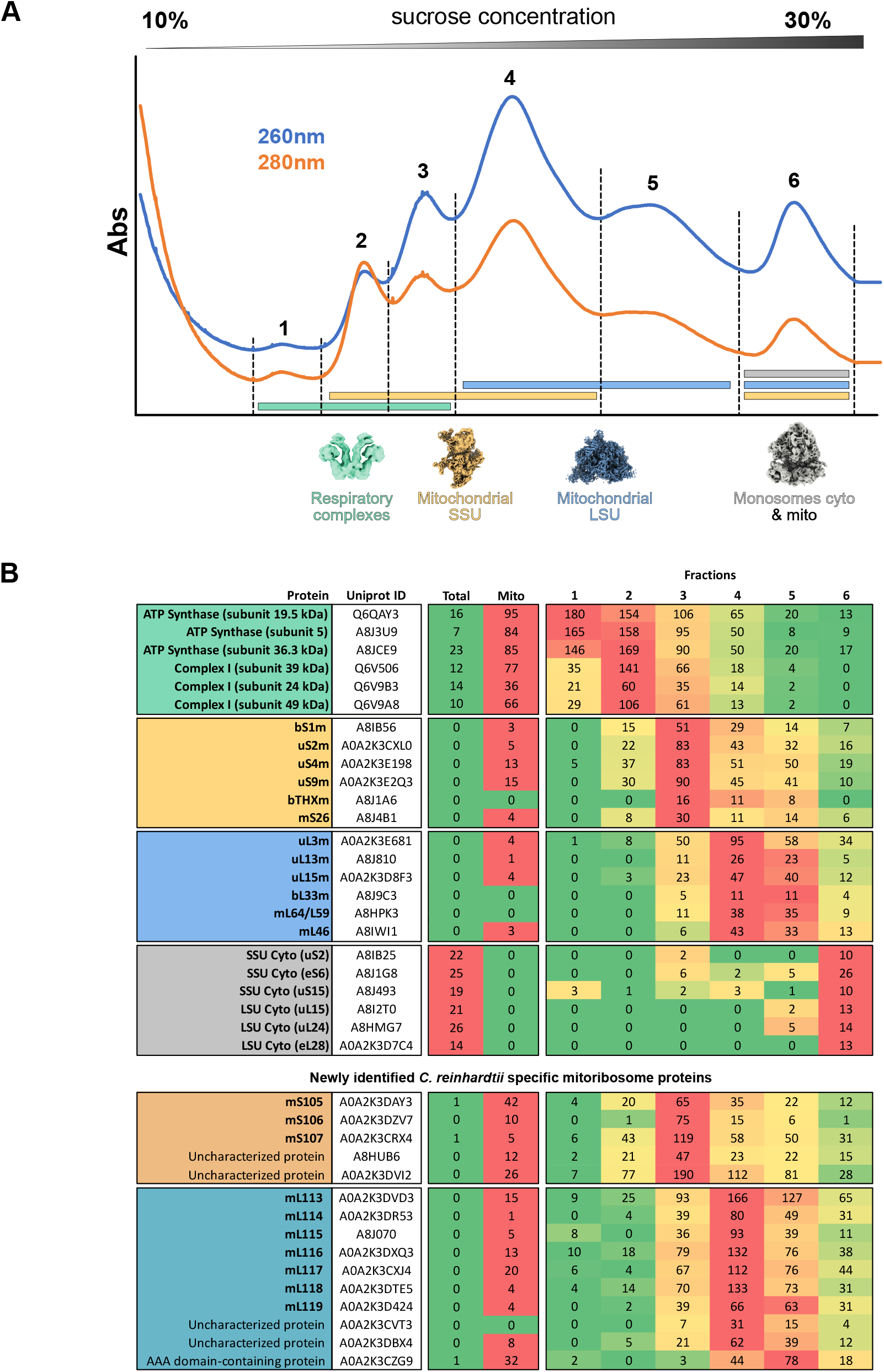
*C. reinhardtii* mitoribosome purification and identification of novel ribosomal components. **A)** Continuous sucrose gradient profile (260 and 280 nm absorbance). 6 peaks were analyzed by proteomics and screened by cryo-EM. Repartition and reconstructions of the complexes identified are shown below the chromatogram. **B)** Summary of the proteomic analysis of the corresponding fractions presented in **A** (full data in Extended), that allowed the identification of novel r-proteins. Total cell (Total) and purified mitochondria (Mito) fraction MS analyses are also shown here. The abundance of the respective proteins is represented as absolute spectra values colored such that higher spectra values are red and lower are green. The upper part of the table presents known components of the complexes identified that were used as references to identify novel components of the LSU and SSU of *C. reinhardtii* mitoribosome. The novel components are presented in the lower part of the table. Proteins that were confirmed by cryo-EM are shown in bold.

**Figure S2.**
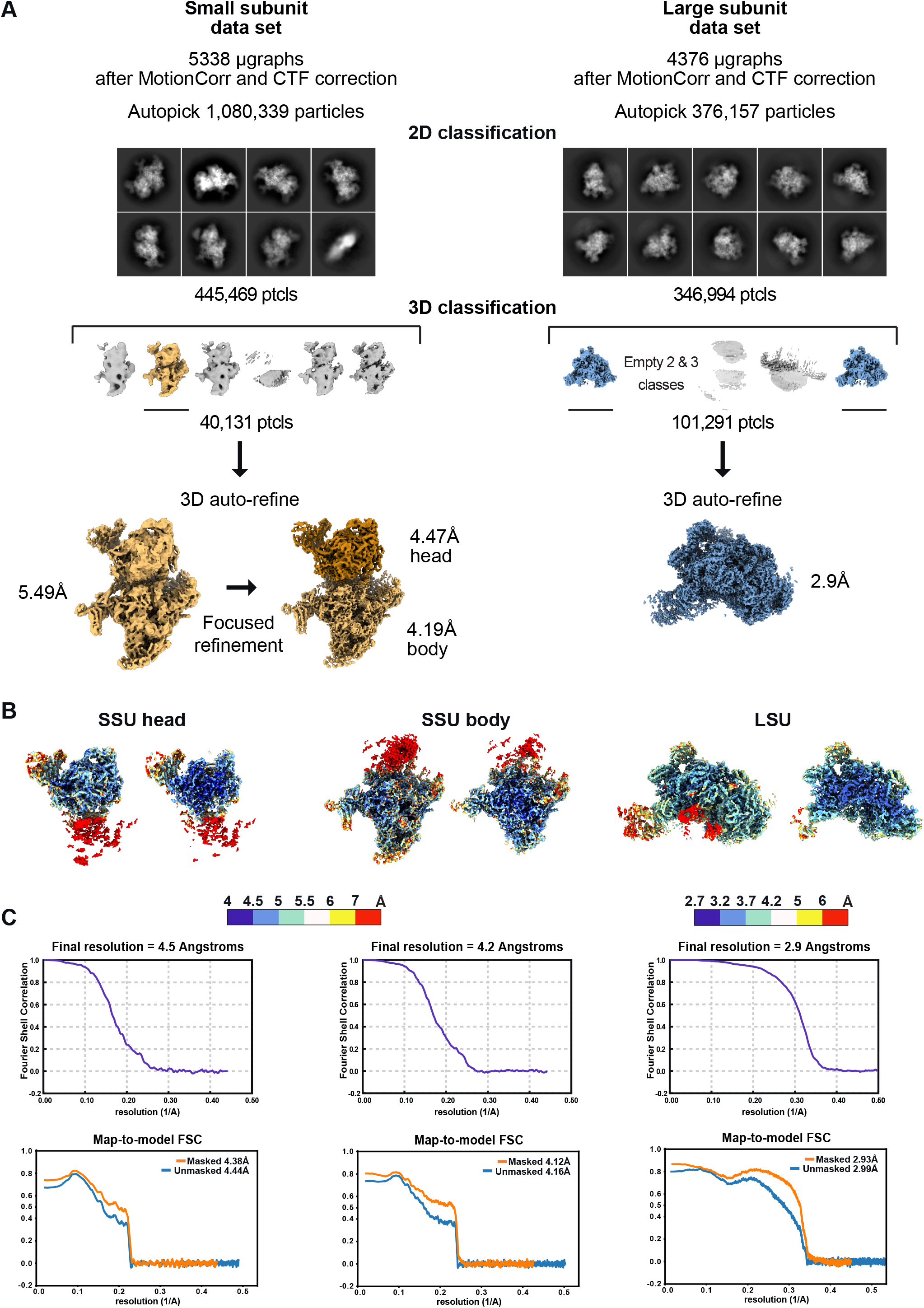
Single-particle data processing workflow. **A)** Graphical summary of the processing workflow described in Methods, with 2D and 3D classes, processing and refinement. **B)** Local resolutions of both the LSU and each focused SSU are presented. The maps are colored by resolution, generated using ResMap (Kucukelbir et al., 2014). Maps are also shown in cut view. **C)** For each reconstructions FSC plots (output from RELION (Zivanov et al., 2018)) are displayed for resolution estimation. Map to model FSC are also shown (output from PHENIX (Liebschner et al., 2019) validation). The maps resolution were calculated at the 0.143 threshold.

**Figure S3.**
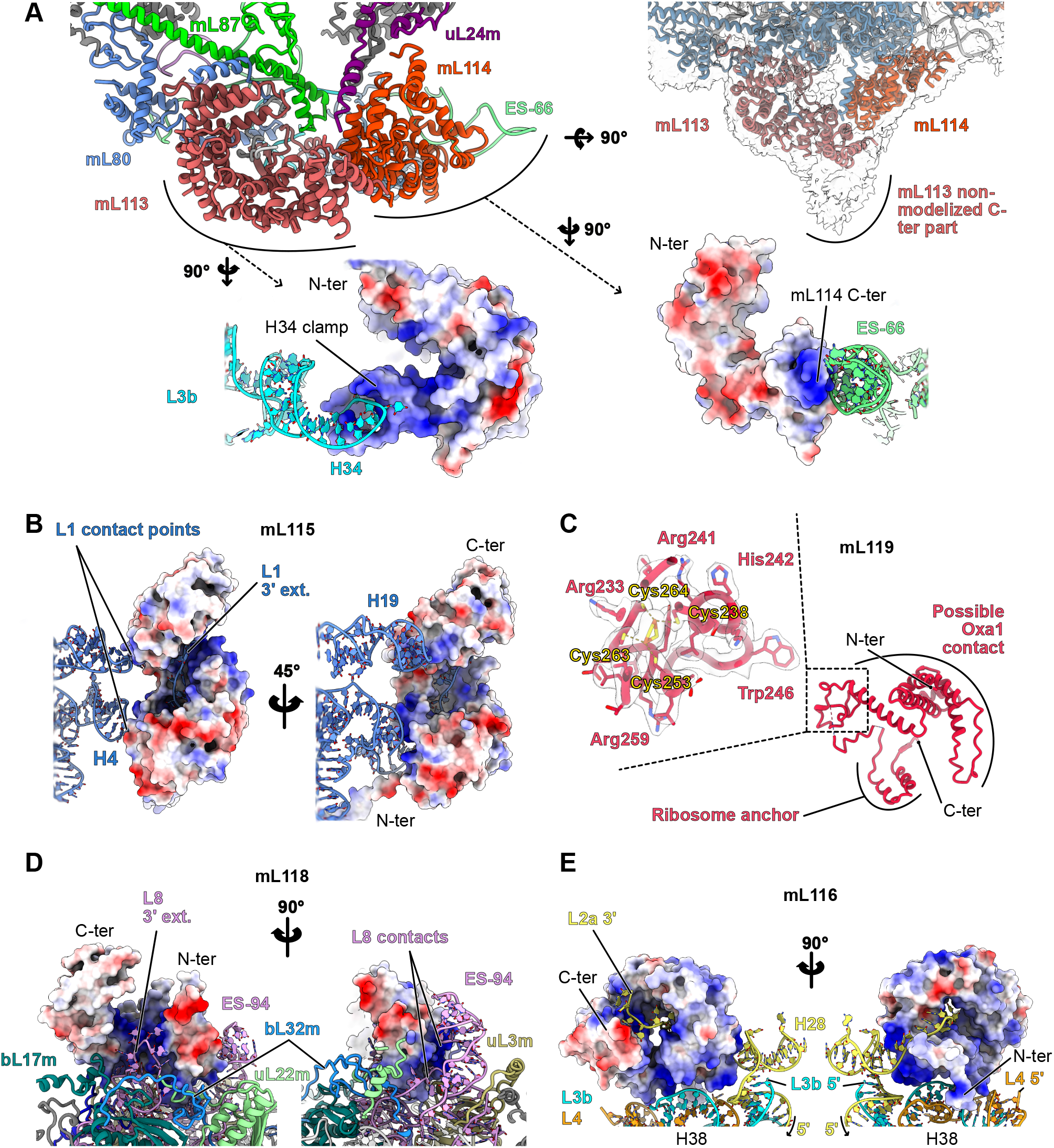
Detailed analyses and views of the specific LSU proteins. Focused views of the different Chlamydomonas-specific r-proteins of the LSU. **A)** View of the mL113 – mL114 area. Both proteins interact together and with the surrounding proteins. The non-modelized C-terminal part of mL113, interacting with the membrane, is shown. It interacts with H34 of the L3b fragment. mL114 interacts with ES-66 of the L7 fragment. Electrostatic coloration (blue coloration corresponds to positive patches and red to negative patches) of the proteins correlate with their rRNA binding site. **B)** Electrostatic coloration of the mL115 protein reveal that its inner groove is mostly positively charged similarly to the two contact points made with H4 and H19. **C)** Detailed view of mL119. Part of the protein coordinate an iron-sulfur cluster *via* four cysteine residues. The model is shown in its density. **D)** Electrostatic coloration of the mL118 protein showing the positive charge of the inner groove and the two contact points made with ES-94. **E)** Similarly mL116 by electrostatic potential revealing the largely positive surfaces both in its inner groove and exterior to interact with rRNAs.

**Figure S4.**
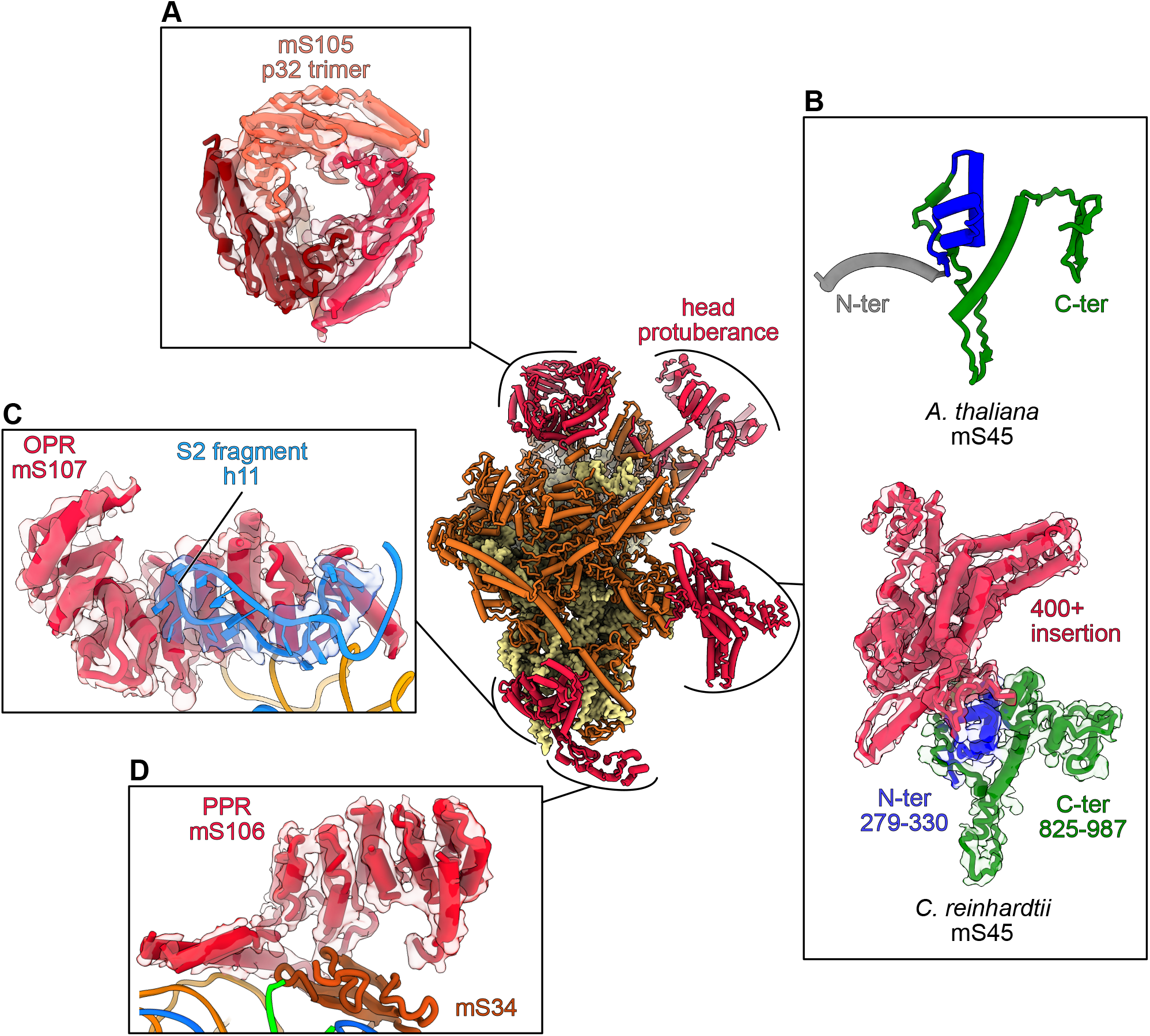
Detailed analyses and views of the specific SSU proteins. The small subunit model is presented from the solvent view which allows to see the major structural features, highlighted in red, of the SSU. Two Y-shaped protuberances are observed on the head and the body. **B)** The body protuberance is formed by a large insertion in the conserved mS45 protein. A comparison between the Arabidopsis and Chlamydomonas mS45 proteins are shown. The head protuberance components, could not be clearly identified and was thus entirely build as poly-alanine. However, several conserved proteins of the SSU head, notably uS3m, uS10m and mS35, have large parts that could not be modelized. These extensions could come together and form this large head protuberance. The foot extension is formed by two helical proteins, one PPR (mS106) (**D**) and one OPR (mS107) (**C**). The PPR occupies a position similar to mS27 in human (Amunts et al., 2015; Greber et al., 2015) and fungi (Itoh et al., 2020), but does not appear to interact with RNA. (**C**) The OPR encapsulates the tip of helix 11. **A)** On the back of the SSU head, the homotrimeric torus-shaped domain formed by three copies of mS105 (p32) is present.

**Figure S5.**
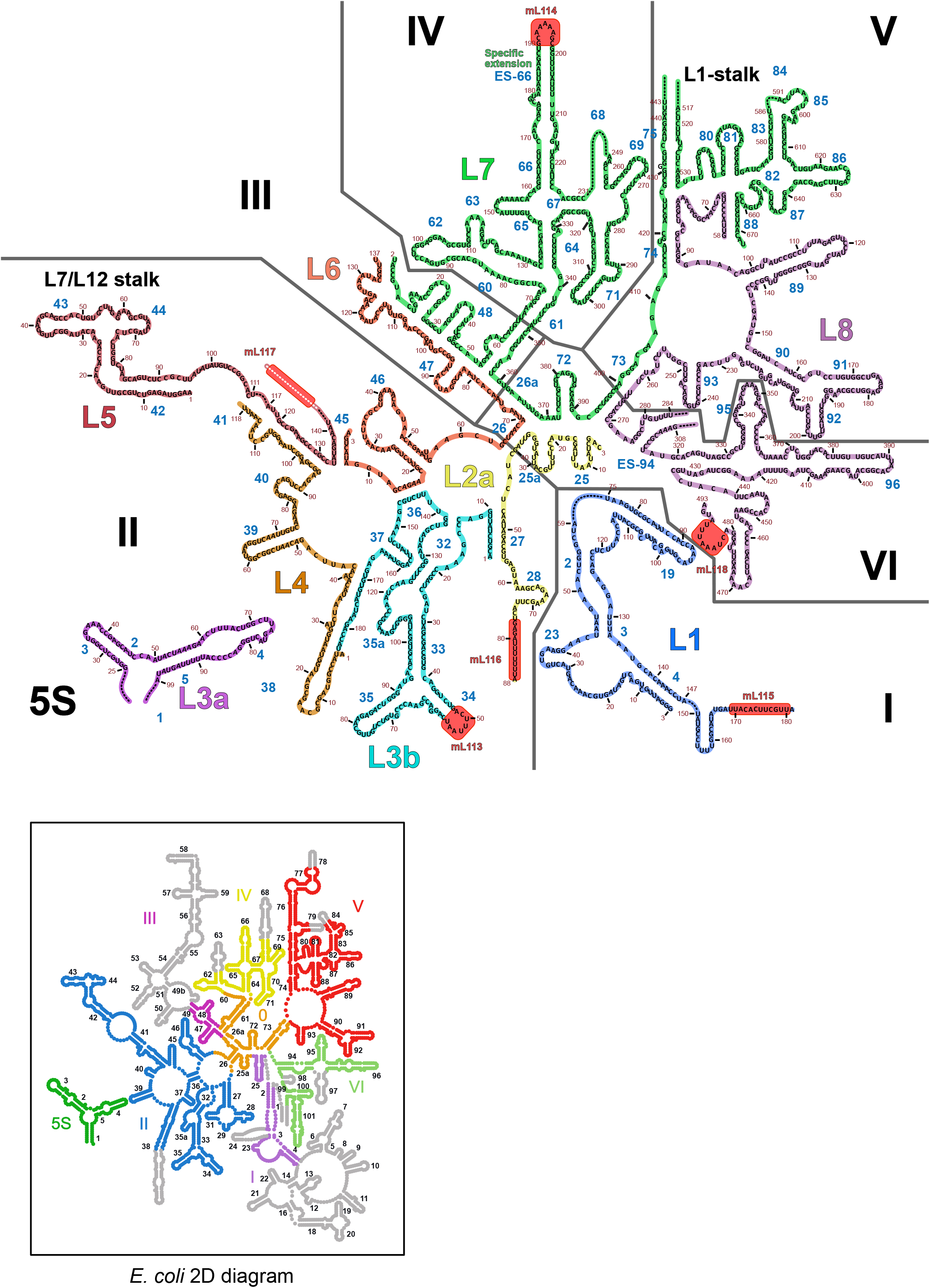
2D diagram of the *C. reinhardtii* LSU rRNAs. 2D representation of the LSU rRNAs of Chlamydomonas mitoribosome. Each 9 rRNA fragments are colored differently and match the color code of Figure 3. The rRNA expansions (ES-94 and ES-66) are indicated on the diagram. RNA portions that could not be modelled are indicated by dark dashed lines, e.g the L1 stalk could not be built due its motion. White dashed lines indicated regions that were built as polyU/A. Contact points of Chlamydomonas-specific r-proteins are also highlighted by red boxes. Simplified secondary structure diagram of the *E. coli* 23S rRNA is also shown in the black frame, this time colored by domain. Helices absent in Chlamydomonas mitoribosome are shown in gray, which highlight the strong reduction of domain I and III. Secondary structure templates were obtained from the RiboVision suite (http://apollo.chemistry.gatech.edu/RiboVision).

**Figure S6.**
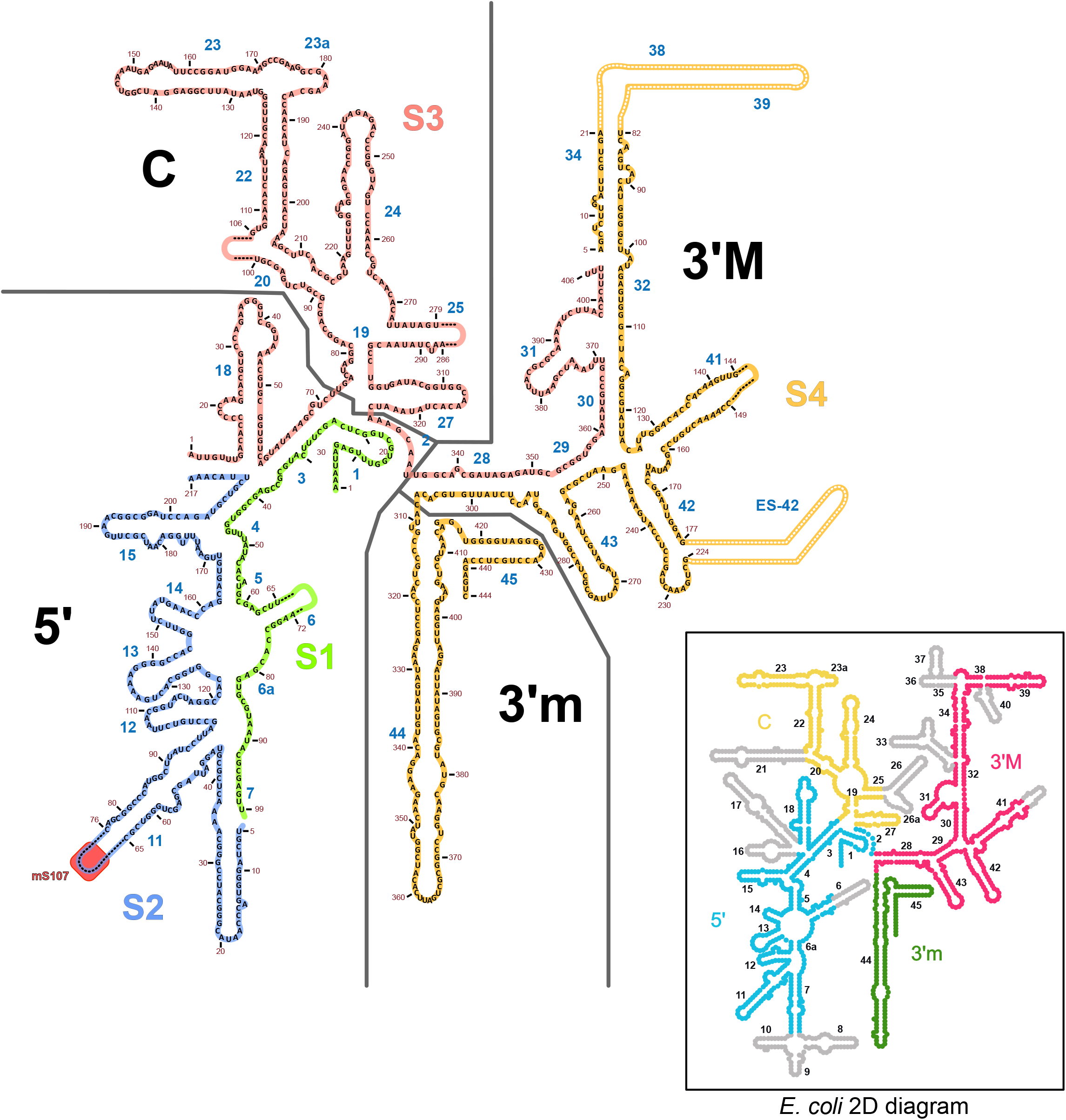
2D diagram of the *C. reinhardtii* SSU rRNAs. 2D representation of the SSU rRNAs of Chlamydomonas mitoribosome. Each 4 rRNA fragments are colored differently and match the color code of Figure 3. The rRNA expansions are indicated on the diagram. Extensions that could not be modelled are indicated by dashed lines. White dashed lines indicated regions that were built as polyU/A. Contact points of Chlamydomonas-specific r-proteins are also highlighted by a red box. Simplified secondary structure diagram of the *E. coli* 16S rRNA is also shown in the black frame, this time colored by domain. Helices absent in Chlamydomonas mitoribosome are shown in gray. Secondary structure templates were obtained from the RiboVision suite (http://apollo.chemistry.gatech.edu/RiboVision).

**Figure S7.**
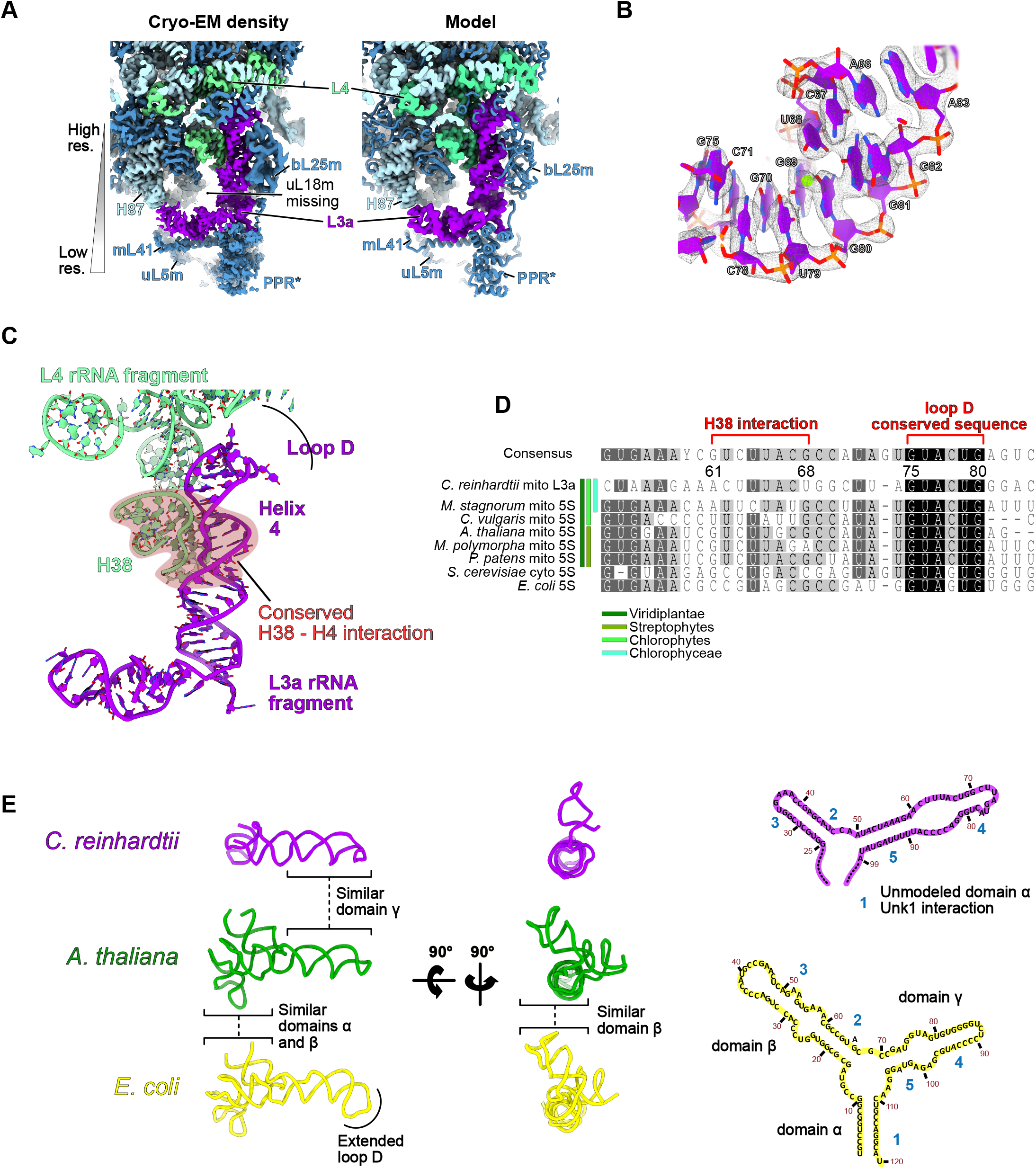
Analysis of the central protuberance, the L3a rRNA fragment is a divergent 5S rRNA. **A)** Comparison of the cryo-EM map of the central protuberance (CP) with the corresponding atomic model. Resolution is lower at the outer extremity of the CP due to the CP general motion. rRNAs are shown in light blue, aquamarine (L4) and purple (L3a) and proteins in blue. For the atomic model, rRNAs are shown in surface and proteins in cartoon representation. Compared to classical CP structures, the uL18 protein is missing, which is normally involved in domain α stabilization. **B)** L3a atomic model in its respective density that allowed identification of the rRNA fragment. **C)** The structurally conserved helix 4 area, interacting with helix 38 of the L7 fragment is highlighted and sequence alignment of the area is shown in **D**. Conservation is observed from bacteria to eukaryotes, especially at the sequence level for the pre-loop D area. The L3a fragment is particularly divergent even with member of the Chlorophytes. **E)** shows structural comparison between bacterial, mitochondrial higher plants and Chlamydomonas 5S rRNAs. Domain γ, especially helix 4 and the D loop are more similar between *A. thaliana* and *C. reinhardtii* compared to bacteria, whereas domain β is completely different in *C. reinhardtii* compared to both bacteria and *A. thaliana.* Chlamydomonas domain β is angled differently which allows the interaction of the final loop of the domain with H87 which is not the case in other ribosomes. The unmodeled domain α of Chlamydomonas L3a most likely interacts with the unidentified alpha helical protein, resembling a PPR protein, PPR*.

**Table S1.**
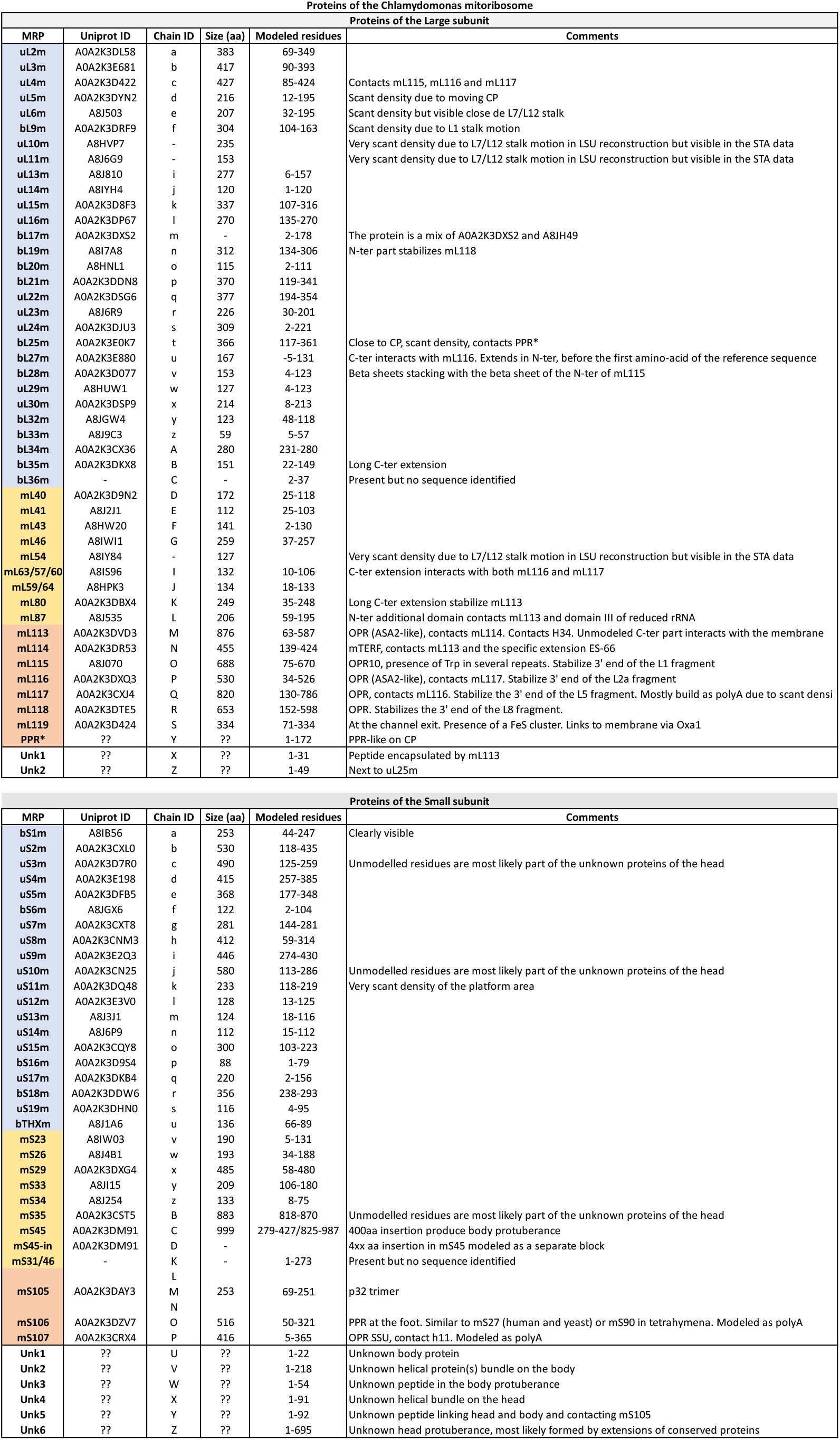
List of proteins identified as constituent *C. reinhardtii* mitoribosome, for the LSU and SSU. List of the r-proteins constituting the *C. reinhardtii* mitoribosome, the table is divided between LSU and SSU r-proteins. The proteins are colored by conservation with the bacterial ribosome (blue) other mitochondrial ribosomes (yellow) or specific to *C. reinhardtii* mitoribosome (red). Due to the L7/L12 stalk motion, proteins uL10m, uL11m and mL54 were not visualized and are presented only in the illustration figures, not the deposited final model. Similarly, bL12m was observed in the subtomogram averaging reconstruction but is not present in the final model. Due to the overall lower resolution of the SSU compared to the LSU, the totality of visible extensions and insertion were modelled as polyA. bS21m, mS37 and mS38 are most likely present but could not be observed in our reconstruction. Full list of proteins identified is provided in Supplementary information.

**Table S2.**
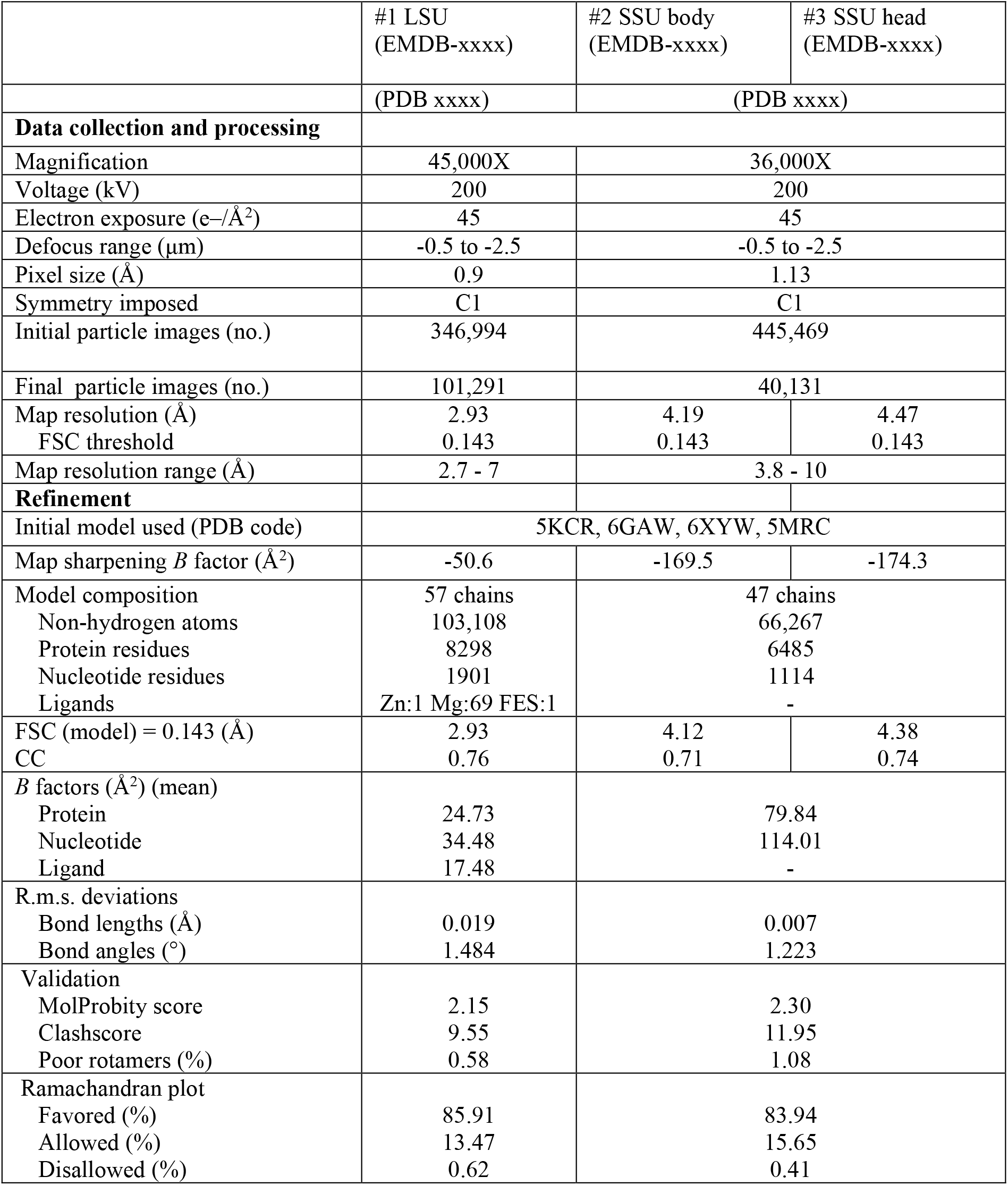
Cryo-EM data collection, refinement and validation statistics.

**Table S3.**
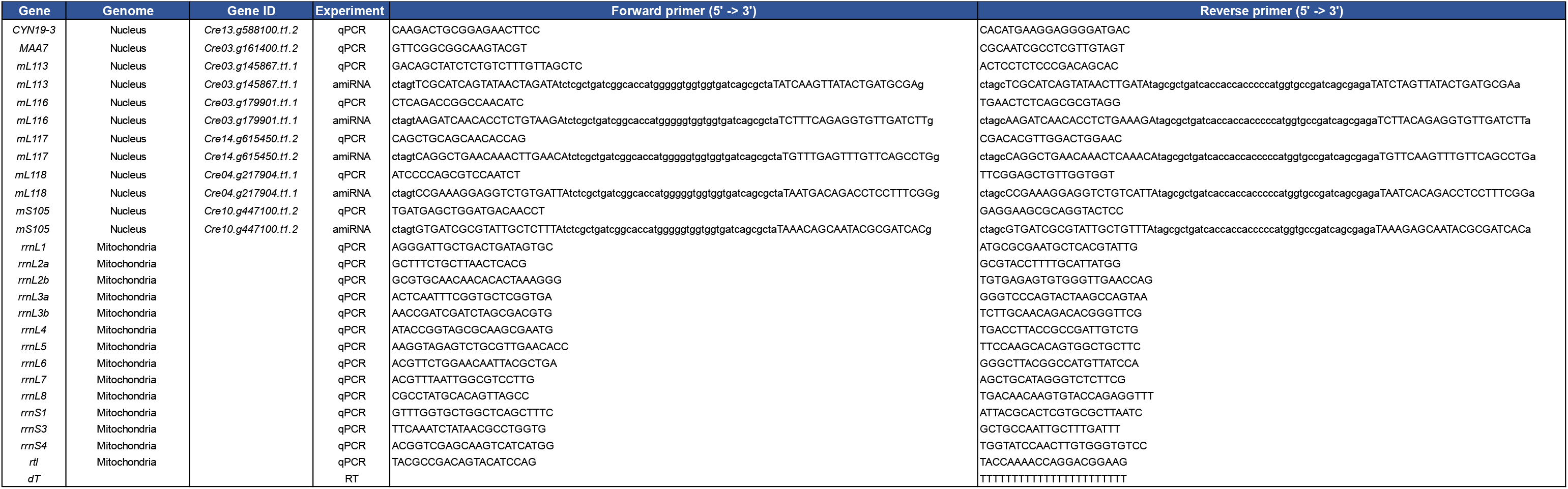
List of oligonucleotides used in this study.

